# Design and experimental characterization of specificity-switching mutational paths of WW domains

**DOI:** 10.64898/2025.12.08.693000

**Authors:** Ahmed Rehan, Eugenio Mauri, Jorge Fernandez-de-Cossio-Diaz, Pierre-Guillaume Brun, Remi Monasson, Marco Ribezzi-Crivellari, Simona Cocco

## Abstract

Specific interactions between proteins and other biomolecules are ubiquitous in cellular processes. How specificity is encoded in the protein sequence and can be modified through a minimal set of concerted mutations is a complex issue. In this work, we focus on the WW protein domain, whose variants specifically bind to different classes of proline-rich peptides. Combining unsupervised learning of homologous WW sequence data with Restricted Boltzmann Machines (RBM) and path-sampling methods, we design mutational paths of putative WW domains interpolating between two natural WW domains with either distinct or similar specificities. Sequences along the designed paths are then experimentally validated with high-throughput in-vitro binding assays against 3 peptides of different classes. The vast majority (93%) of intermediate sequences along the designed paths are responsive to the initial or/and final peptides. On the contrary, domains along scrambled paths, in which the same mutations are introduced in random order are not functional, emphasizing how successful design crucially depends on the ability to model epistatic interactions. Switch in specificity between classes I and IV, whose representative peptides bind to different pockets on the WW domain takes place through intermediates displaying some level of binding cross-reactivity with the tested peptides, contrary to the transition from class I to II, which are associated with the same binding pocket. Lastly, we show that the RBM paths share a high identity with internal nodes obtained from ancestral sequence reconstruction based on the seed WW domains.

**Significance Statement:** Generative machine-learning models are nowadays used to design new protein sequences with desired functions. Here, we address a more demanding task: designing a full mutational path connecting two natural proteins with different binding specificities. We illustrate this problem with WW domains, a small protein unit capable of recognizing distinct classes of proline-rich peptides. We experimentally verify that most of the intermediate sequences along the designed path are functional and respond to the initial or/and final peptides. The designed sequences share significant homology with the sequences obtained as internal nodes of phylogenetic trees through ancestral sequence reconstruction.

## 1 Introduction

As living organisms evolve, the proteins they produce may need to adapt and specialize to a new task while preserving their fundamental biological function, such as catalyzing certain classes of reactions or binding to specific targets [1]. An illustration of this flexibility is provided by hemoglobin and myoglobin: these proteins share a common ancestor and the conserved mechanism of binding oxygen to the heme group, however hemoglobin transports oxygen from the lungs to the tissues, and myoglobin stores oxygen for use during muscle contraction [2]. Another interesting example is the WW domain, a family of small protein domains (30-40 amino acids long), whose name comes from the presence of two strongly conserved tryptophan residues [3]. Found in proteins across multiple species, WW domains are essential to key cellular pathways, such as the Hippo pathway, which controls cell proliferation, apoptosis, and organ size [4]. WW domains are widely distributed among proteins and are commonly found both in the cytoplasm and in the nucleus [5]. Following their initial identification, these domains became a primary focus of scientific inquiry due to their involvement in signaling complexes linked to a range of human disorders, including some types of cancers, Alzheimer’s disease, muscular dystrophy, and cardiovascular disorders [6].

The involvement of mutations or dysregulation of WW domain-containing proteins in the pathogenesis of various diseases, has been well documented. For instance, aberrant activity of the YAP/TAZ transcriptional regulators, which is mediated by their WW domains, is a hallmark of many cancers due to its role in promoting unchecked cell growth [7]. Similarly, mutations in the WW domain of Pin1 are involved in the emergence of Alzheimer’s disease [8], while NEDD4-2 are associated with Liddle syndrome, which disrupts sodium channel regulation and is a risk factor for hypertension [9]. These examples illustrate the significance of WW domains in health and disease, establishing them as crucial targets for therapeutic intervention.

Most WW domains bind proline-rich peptides [10]. However, they differ in the specific amino-acid motifs they recognize, defining three broad specificity classes (I, II/III, IV) [11, 6, 12], which determine the biological pathways their interactions affect. Two examples of WW domains present in the human proteome are the yes-associated protein 1 (YAP1) and the Peptidyl-prolyl cis-trans isomerase NIMA interacting 1 (Pin1). While the former acts as a transcriptional co-activator of genes involved in cellular proliferation (with a major role in oncogenic activity [13]), the latter isomerizes only phospho-Serine/Threonine-Proline motifs and has many roles, among which cell cycle regulation. Rational design of WW domains has been proposed to change their specificity and recognize mutated ligands. As an example, in Liddle’s syndrome, the PPXY motif of a sodium channel subunit, *β* EnaC is mutated, which prevents sodium channel degradation by Nedd4; changing specificity of the WW domain from class I to II/III would restore recognition [14].

In addition to their clinical relevance, WW domains serve as valuable models in structural biology and protein evolution due to their small size, conserved structure, and diverse binding partners. Their modular nature and capacity to mediate specific interactions provide insights into protein network dynamics and evolutionary adaptation [15, 11, 6, 12, 16]. In this context, a natural question is to understand how homologous proteins with different ligand specificities evolved from ancestral, and possibly promiscuous proteins [17], while remaining functional at all stages. This question dates back to the seminal works of J. Maynard Smith, who described evolution as a word game in which one can change one letter at a time while keeping meaningful intermediate words, as illustrated by the path “word*→*wore*→*gore*→*gone*→*gene” [18]. Ancestral sequence reconstruction (ASR), by constructing putative intermediates linking the diverse modern proteins, is a practical way to address this problem [19]. However, bridging the gap between the theoretical and conceptual study of evolution through paths [20] and realistic fitness landscapes is difficult [21, 22].

From an experimental point of view, high-throughput experiments allow for building mutational paths between phenotypically distinct proteins separated by *D* mutations [22, 23, 21, 24, 25], when *D* is small enough. Examples include Ref. [25], in which mutational paths in a region of 4 residues increasing antibiotic resistance in *β*-lactamase were investigated, and Ref. [21], where all 2^13^ direct paths involving 13 residues allowing to change the fluorescence properties of a protein, were tested. Let us also mention Ref. [26] where all 20^4^ paths over a 4-residue region in the protein GB1 were characterized and Ref. [27], where about 1000 paths in the fitness landscape of transcription factors for short (9 nucleotides) DNA sequences were tested and characterized. When the number *D* of mutations separating the two proteins becomes large, the space of possible sequences cannot be explored in an exhaustive manner any longer.

Evolutionary methods such as ASR can help to isolate sets of important mutations involved in the switch of affinity in protein systems [28, 29], although the phenotypic characterization of the mutations remains essentially experimental. Computational models approximating the genotype-phenotype mapping are necessary to prune this huge space and retain only good putative paths leading to the change of function.

Recently, we proposed such a computational approach to design mutational paths [30]. Briefly speaking, the approach relies on two ingredients. First, we use an unsupervised machine-learning architecture, trained from homologous sequence data, to model the global fitness landscape of the protein family of interest, hereafter WW domains. Various generative models were recently proposed based on deep architectures [31, 32], e.g. diffusion processes [33]. Simpler unsupervised architectures such as Boltzmann Machines (BM) or Variational Auto-Encoders have been shown to be good generative models, with the ability to design novel proteins with functionalities comparable to natural ones [16, 12, 34, 35, 36], while being easier to interpret. Hereafter, we use Restricted Boltzmann Machines (RBM), which are bipartite probabilistic graphical models learning relevant amino-acid motifs with key role for structure stability and function [37] and able to design new functional proteins [38] or other biomolecules [39, 40]. Second, we sample the space of mutational paths, defined as chains of sequences that (1) have high scores according to the RBM generative model and (2) differ from the previous and next ones along the path by a small (and controllable) number of mutations. At their extremities the chains are anchored by two sequences of interest, hereafter two natural WW domains with distinct specificities.

This path design approach was shown to successfully generate high-quality mutational paths of in silico (lattice-based) proteins, for which the ground-truth was known [30]. In addition, we showed it could be applied to the WW domain family, using as training data the multiple-sequence alignment from the PFAM/InterPro database (PFAM ID: PF00397). Intermediate configurations had high scores according to the trained model and to computational methods based on protein structure information, such as AlphaFold and ProteinMPNN [41, 42].

In this paper, we go a step further in designing mutational paths for switching specificity in WW domains. We present a high-throughput fully in-vitro binding assay that allows us to simultaneously evaluate the activity of the designed WW domain paths against multiple peptides attached to different specificity classes. These experimental results not only validate the computational approach but also allow us to study in great details where and how the switching of specificity takes place along the path. Two key questions are: i) is the transition happening through an abrupt switch of specificity (punctuated equilibria, corresponding to distinct specificities) or through gradual changes (with intermediate that exhibit mixed or intermediate specificity)?; ii) is the switch associated with a bottleneck in sequence space and very few possible paths? An abrupt switch with a bottleneck has been observed in [21] for the fluorescent protein. Lastly, we show that the RBM designed paths share some similarity with the internal nodes of phylogenetic trees inferred from representative WW domains sequences (the PFAM seed), enriching the number and quality of candidate domains and potential evolutionary scenarios compared to what is achievable by traditional protocols such as ASR.

## 2 Computational models for path generation and experimental verification

Our pipeline is described in Figure 1. We consider the space of putative WW domains (Figure 1A), whose sequences contain a variable portion of *L* = 31 (Figure 1B) amino acids plus appropriate constant flanking regions or ’spacers’ (Figure 1D). The probabilistic model for the variable part is provided by a two-layer unsupervised neural network, called Restricted Boltzmann Machine (RBM), trained on homologous natural sequence data, see Figures 1B,C (Methods 5.1 and [43, 30]). RBM learns a distribution over the set of all possible sequences, which allows us to score putative WW sequences, and, in turn, to generate new sequences with high scores.

**Figure 1:**
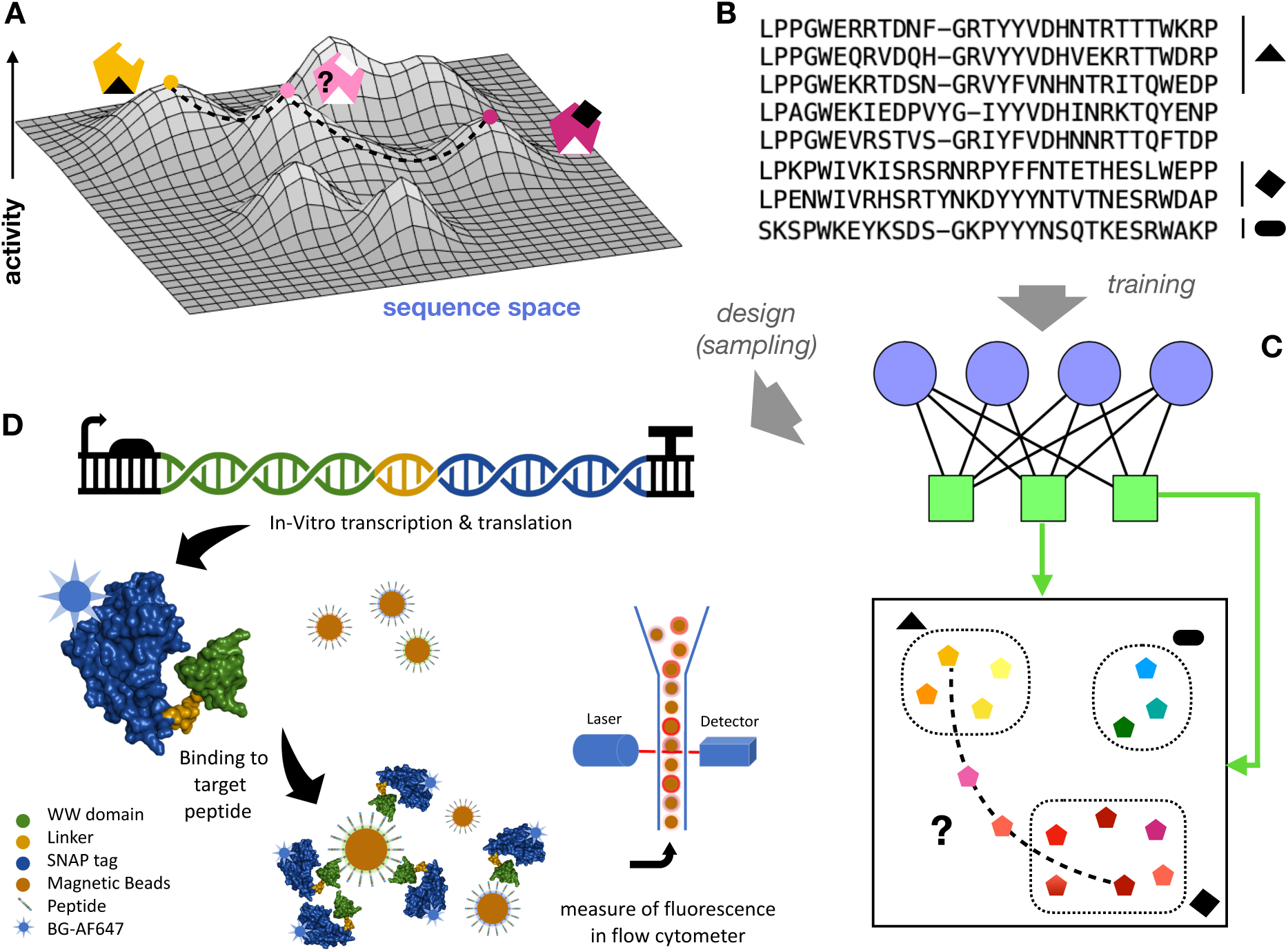
Experimental and computational pipeline for designing WW domains of variable specificities. **A.** Sketched of the activity vs. sequence landscape of WW domains. Highly active domains may bind to different peptides (black shapes) and are shown with different colours. **B.** Homologous sequences define the training data for our restricted Boltzmann machine (RBM) model. The binding specificities of some sequences are annotated, while others are not available. **C.** After training, the latent variables of the RBM define low-dimensional projections that identify clusters of sequences sharing the same specificity. Sequences along paths may cross regions deprived of natural sequences, of unknown specificity. **D.** Sketch of the experiment. <u>In vitro</u> transcription and translation of our construct result in the expression of a fusion protein including a WW domain (green), a linker (yellow), and the SNAP tag (blue), labelled with BG-AF647 dye. The WW domain of the fusion protein may bind to peptide-coated magnetic beads (brown). The fluorescence intensity on the bead surfaces is then measured using flow cytometry, assessing the strength of the interaction between the labelled protein and its target peptide.

Natural WW domains are broadly classified according to three specificity classes (I, II/III, IV) [11, 6, 12]. We choose peptides representative of each class to assess the binding specificity of the designed variants (Table 1). Some latent variables inferred by the RBM have been shown to be informative about the specificity classes [43, 30]. More precisely, high-dimensional sequences cluster in the 2-dimensional plane defined by the inputs attached to two particular latent units, *I*_1_ and *I*_2_ (see Ref. [43], Methods Section 5.2, Figure 2A and Suppl. Figures 8 and 9). Experimentally tested sequences reported in the literature appear to consistently belong to the same clusters depending on their specificities. This clustering property is sketched in Figure 1C.

**Figure 2:**
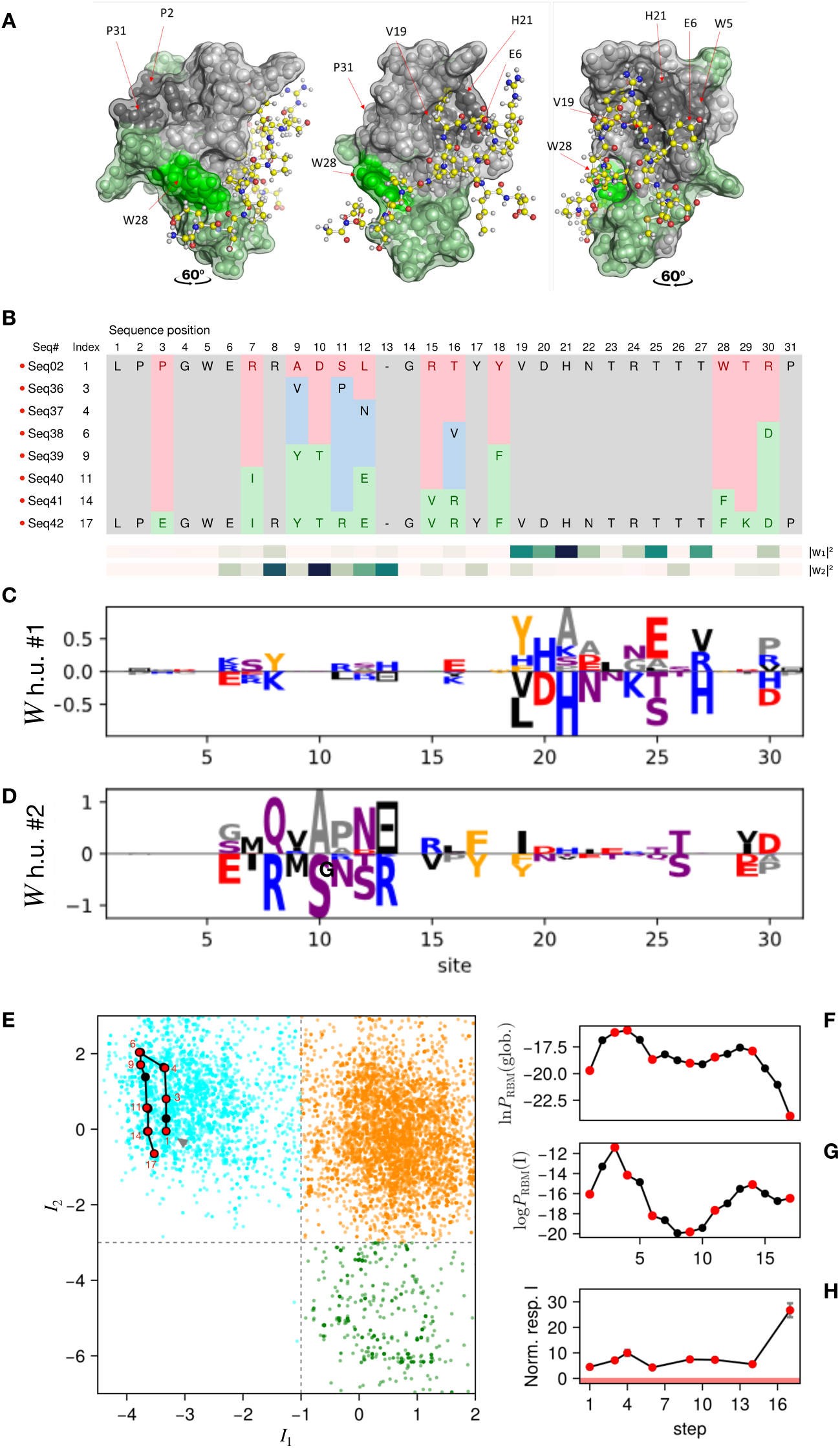
Path within class I WW domains. **A.** Wild-type WW domain NP595793.1 in complex with its peptide (Ligand) shown from both sides, predicted by Colabfold. Amino-acid residues that change along the path are highlighted in green. **B.** Alignment of the tested sequences showing the mutated amino acids. Sequence identity number (Seq#) and index (Index) along the path are specified. **C, D.** Logo representation of the weight vectors attached to the two RBM latent units most correlated with the specificity classes. **E.** Projection of the path on the latent weights, clustering sequences according to their binding specificities. The specificity thresholds indicated in the figure (Class I: *I*_1_ *<* −1 and *I*_2_ *>* −3; Class II/III: *I*_1_ *>* −1 and *I*_2_ *>* −3; Class IV: *I*_1_ *>* −1 and *I*_2_ *<*-3) are informed by previous specificity assays on natural sequences, reported as crosses in Suppl. Figure 7 [12, 11] (see Figure 3 in [43] and Figure 2a in [30]). **F, G.** Global and Class I specific RBM scores for the sequences in the path. **H.** Experimental binding responses of the tested sequences. The red band, defined by the experimental noise limit, denotes the non-binding zone.

**Table 1:**
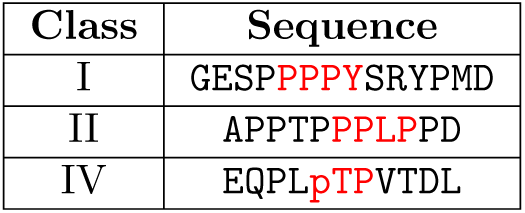
Peptides used for the experimental validation of mutational paths in WW domains. Highlighted in red are residues that bind to the WW domain. The phosphorylated Threonine in the peptide of class IV is indicated by `pT`.

We learn global RBM models, trained on all available WW data irrespectively of their binding specificity, and class-specific models, trained from sequences attached to one class only, see Methods 5.2; notice that the attribution of sequences to one of the three functional classes for training is mostly based on the 2-dimensional clustering mentioned above and not on experimental characterization, and is therefore subject to errors. Class-specific models are, by construction, informative about the binding specificity of the WW sequences but are generally of lesser quality than the global model, due to the smaller number of WW data, in particular for class IV. As a consequence, they cannot be used to generate <u>de novo</u> high-quality sequences from one class or another, but rather to score the specificity of sequences generated with the global model.

We use path-sampling algorithms, developed in [30], to design paths of sequences fulfilling the following conditions:

- the initial and final sequences are natural WW domains, possibly with different specificities, <u>e.g.</u> triangle and diamond in the sketch of Figure 1, and identity (fraction of identical residues) ranging from 30% to 50%, which are typical values within a homologous family.
- Intermediate sequences along the path have high scores (probabilities) according to the RBM global model. Amino acids in the intermediate sequences can take any value.
- The number of mutations from one sequence to the next along the path is small, typically one or two. The length of the designed path is typically of 1.5 times the hamming distance between the initial and final sequences.

To sample paths we have employed a Monte Carlo algorithm, which randomly draws paths according to the RBM probability distribution (Methods 5.3). We have designed four batches of sequences, listed in Appendix B together with the parameters used for the models. Intermediate sequences along the paths are scored according to the global RBM model, and to class-specific models to predict their specificities. In our sampled paths amino acids can ”transiently” mutate into third-party values (underlined in blue in Figure 2A). Transient mutations can yield higher fitness scores along the evolutionary path, particularly when epistatic interactions are strong and compensatory mutations are expected [44].

To test the sequences generated by the model in high-throughput, we developed a fully *in-vitro*, rapid and accurate assay: “fluorescence-Activated Bead Counting (fABC)”. The assay assesses interactions between proteins and target molecules immobilized on magnetic beads, see Figure 1D. In the present case our targets are proline-rich peptides specific of the three different subclasses of WW domains (I, II/III, IV). Briefly, synthetic genes coding for SNAP-tagged WW domains[45] are expressed using cell-free *in-vitro* Transcription and Translation (IVTT) [46]. The SNAP-tag is leveraged for the fluorescent labelling of the WW domains using a benzyl-guanine modified fluorescent dye. Fluorescent binders are then mixed with magnetic beads coated with the target peptide and allowed to equilibrate. Binding activity is then measured by quantifying bead fluorescence on a flow cytometry instrument[47]. We demonstrate how, by using spectrally distinct fluorescently-labelled DNA linkers to coat the magnetic beads, we are able to multiplex different targets within a single run, increasing speed and reducing the cost of the assay. This method provides a robust and versatile platform for studying protein/protein or protein/peptide interactions and sequence/function relationship.

## 3 Results

We present and discuss hereafter a path connecting two natural WWs of type I, three paths connecting natural WWs of types I and IV, two paths connecting natural WWs of types I and II/III. Along each designed path, we pick up the sequences at regular intervals for experimental measurements of their binding affinities to ligands of classes I, II/III and IV. The density of experimentally assessed sequences is higher close to specificity switching. To test the presence of epistatic effects we compare our designed paths with scrambled paths, in which the same list of mutations present in the paths generated by the model are introduced in a different order, hence breaking epistatic interactions with the background sequence. A complete list of the designed paths and tested sequences is provided in Suppl. Tables B.

### 3.1 Assessment of natural WW activity levels

Several WW domains perform their function in vivo either as multi-domains or as part of a complex architecture [48, 49] and may require additional activating, post-translational modifications. Our in-vitro assay is based on the expression of individual WW domains and, as a consequence it is possible that binding activity may be lost. To explore and avert these issues, we tested a set of 28 wild-type sequences taken from the homologous sequence data (12 Class I, 15 Class II/III, 1 Class IV) against three peptides, representative of the targets of the three classes. Out of these sequences, a total of 8 did not give any response in our assay (2 classified as Class I and 6 classified as Class II/III). This is in particular the case of the YAP1 WW domain that we considered in [30]. Half of the remaining sequences gave a clear and specific response for the peptides associated to the classes (8 sequences for Class I, 5 for Class II, and 1 for Class IV). Two sequences showed cross-specificity to peptides associated to Classes I and II. In the following we discuss paths starting and ending at sequences with clear and specific responses (Appendix

### 3.2 Path within class I WW domains

We start by testing a path between two Class I wild-types (NP595793.1 Pub3 and 2OPA-7 NEDD4). Figure 2A shows one of these natural WW domains in complex with the cognate peptide. We show in Figure 2B the sequences of the two natural WWs of type I we considered, as well as the six tested intermediate sequences, sampled by the RBM-path algorithm connecting the two wild-types that were experimentally tested. The full sampled path is given in Suppl. Fig 11). Residues that change along the path are colored. As shown on the structures in Figure 2A, a large fraction (*↑* 40%) of amino acids are mutated along the path. In Figure 2C,D we show the weights attached to the two specificity-related latent units of the RBM (*I*_1_, *I*_2_). Projections of the sequences in the path are consistently confined within the cluster attached to class I (Figure 2E). Most mutated amino acids are not proximal with the ligand, and not located on the sites with large entries in the RBM-weight attached to the latent units detecting class I specificity (*I*_2_) (Figure 2C, in agreement with the fact that the specificity remains unchanged along the path. Twelve positions in total are altered across the path, four of which (9, 11, 12, 16) sample an intermediate residue, allowing to improve the global score of the artificial intermediate proteins with respect to the initial and final natural wild-type.

Figure 2H shows the measurement of the binding to a class I peptide of the tested sequences in the path. All tested sequences show specificity responses much above noise level. The global RBM scores of the sequences along the path and the class I specific RBM score are shown in Figure 2F,G.

Almost all artificial sequences along the path (Figure 2F) have global RBM scores larger than the anchoring wild-type domains. Sequence sampling is, indeed, biased to generate large RBM-score sequences through the use of a temperature equal to 1*/*3 (see Methods 5.3), proven to be effective in [34]. A general discussion and comparison of the distributions of scores of designed and natural sequences is given in Section 3.5. Interestingly, designed sequences with the largest scores, in particular the second and the third sampled ones, have larger responses than the initial natural WW.

The presence of indirect (transient) mutations (9, 11, 12, 16) improves the global score. In particular, several pairs involving site 9 (9-13, 9-14, 9-16) have a large epistatic score according to the RBM model, and correspond to contacts on the 3D fold, see Suppl. Figure 2. Indirect mutations may be therefore important for stabilizing the designed WWs along the path by exploiting epistatic interactions. As an illustration, the transient mutation A9V appearing on the second tested WW (Seq. 36) increases the stability via a favorable contact interaction, as seen from the value of the Miyazawa-Jernigan energy [50] between amino acids V9 and L12. This interaction is lost on the third tested sequence (Seq. 37) due to the mutation L12N compensated by the mutation T16V through a favorable contact interaction between amino acids V9 and V16. The Class I natural sequence anchoring the path (Seq. 42) has the smallest global score according to the RBM, but a good Class I score and a very good binding response, illustrating the difference between the global score, a proxy for stability, and the specificity scores. In conclusion, the experimental probing of a designed path connecting two Class I wild types allows us to assess the binding response of the designed sequences along the path and to validate the RBM model predictions, in particular the epistatic interactions and the global and class-specific scores.

### 3.3 Paths from Class I to Class IV

We now consider a path designed to interpolate between two different specificity classes, I (wild-type NP595793.1 Pub3) and IV (wild-type 2LB3A PIN1). As shown in Figure 3A and previously reported [51], ligands to Class IV domains bind to a different pocket compared to their Class I counterparts. This alternative binding mode requires a longer loop between the *β*_1_ and *β*_2_ strands. Position 13 carries a gap in the Class I WW domain, and amino acid S in the Class IV natural domain (Figure 3B). The existence of two binding pockets may favor switching specificity and binding cross-ractivity as was hypothesized in

**Figure 3:**
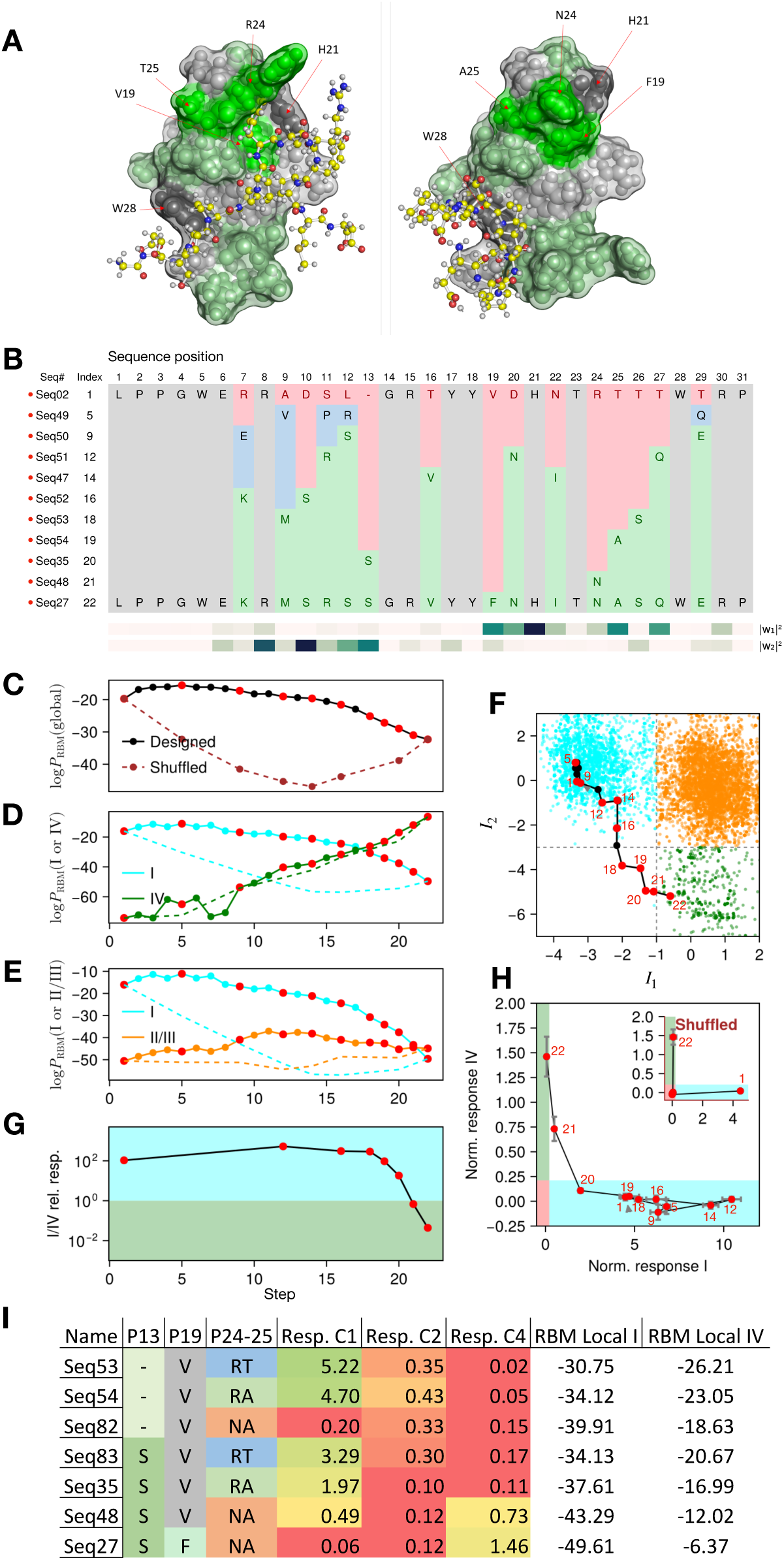
Path from Class I to Class IV. **A.** Structures of the Class I natural WW domain NP595793.1 in complex with Class I peptide (Left) and Class IV natural WW domain 1I8G (Pin1) in complex with Class IV peptide (Right), predicted by Colabfold. Amino-acid residues that change along the path are highlighted in green. **B.** Alignment of tested sequences along the path showing mutated amino acids. Sequence identity number (Seq#) and index (Index) along the path are specified. **C,D,E.** Scores of sequences in the path predicted by the global (top), Class I or IV (middle) and Class I or II/III (bottom) RBM models. **F.** Two-dimensional projection of the path in the plane of specificity-related latent units. **G**. Relative binding responses to Class I and IV peptides of sequences along the path. **H.** Normalizes experimental binding responses to Class I and IV peptides. The colored bands, defined by the experimental noise limit, denotes the non-binding zone. **I.** Responses of Seq.53, 54, 35, 48, 27 compared with Seq.82, 83 differing by mutations on residues 13 (carrying a gap for a shorter *β*_1_ − *β*_2_ loop or an S), and class I specificity-determining positions 24-25.

Class IV poses challenges from the computational point of view: the number of natural sequences in the cluster associated to Class IV is small compared to the other specificity classes, see green dots in Figure 2E. The number of experimentally validated Class IV sequences is even smaller ([11, 12] and Figure 3 in [43]). Due to the lack of training data, we expect RBM models to be less predictive for Class IV sequences.

Figure 3B shows the list of experimentally tested sequences obtained by subsampling the full path (sampled sequences are represented by red dots; the full path is given in Suppl. Fig 12). About 50% of the protein sequence is modified along the path. Most of the mutated residues are within or close to the two binding pockets, which can be recognized based on the large norm of the RBM weights (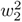 and 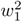 in Figure 3B) of the two specificity-related latent units. The importance of the residues in positions 8 and 13 and their changes in the path to achieve Class IV specificity agree with the weights (previously shown in Figure 2B) attached to unit 1 (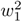 in Figure 4A). On the contrary, sites 19, 24 and 25 carry large entries in the weight (Figure 2C) associated to latent unit 2 (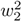 in Figure 3B) [43, 30]. Positions 7, 9, 11, 12, 29 carry indirect mutations in the path (shown in blue in Figure 3A). The global RBM score (Figure 3C) is slightly smaller near the end of the path than at the beginning, perhaps due to the difficulty of properly modeling Class IV-like sequences or to a larger stability of Class I molecules with respect to the other classes, as shown by the MSA alignment in Suppl. Figure 3. We observe the expected trends for the specific RBM models: Classes I and IV have, respectively, lower and higher scores at the end of the path than at the beginning (Figure 3D). The two scores cross after the 6th sampled sequence in the path (Seq. 52). The path then goes, in the low-dimensional projection, across a region devoid of natural sequences (Figure 3E). Interestingly, in this region, the Class II/III specific RBM score gets closer its Class I counterpart (Figure 3E). Experiments confirm that sequences in this region keep a significant activity towards both Class I and Class II ligands (Suppl. Table 3). Furthermore, the last designed sequence along the transition path (WW150) shows cross-reactivity to Class I and IV ligands, consistently with our argument given in [30] (Figure 3H). Taken altogether, these results points towards a substantial level of promiscuity [30] along the path.

**Figure 4:**
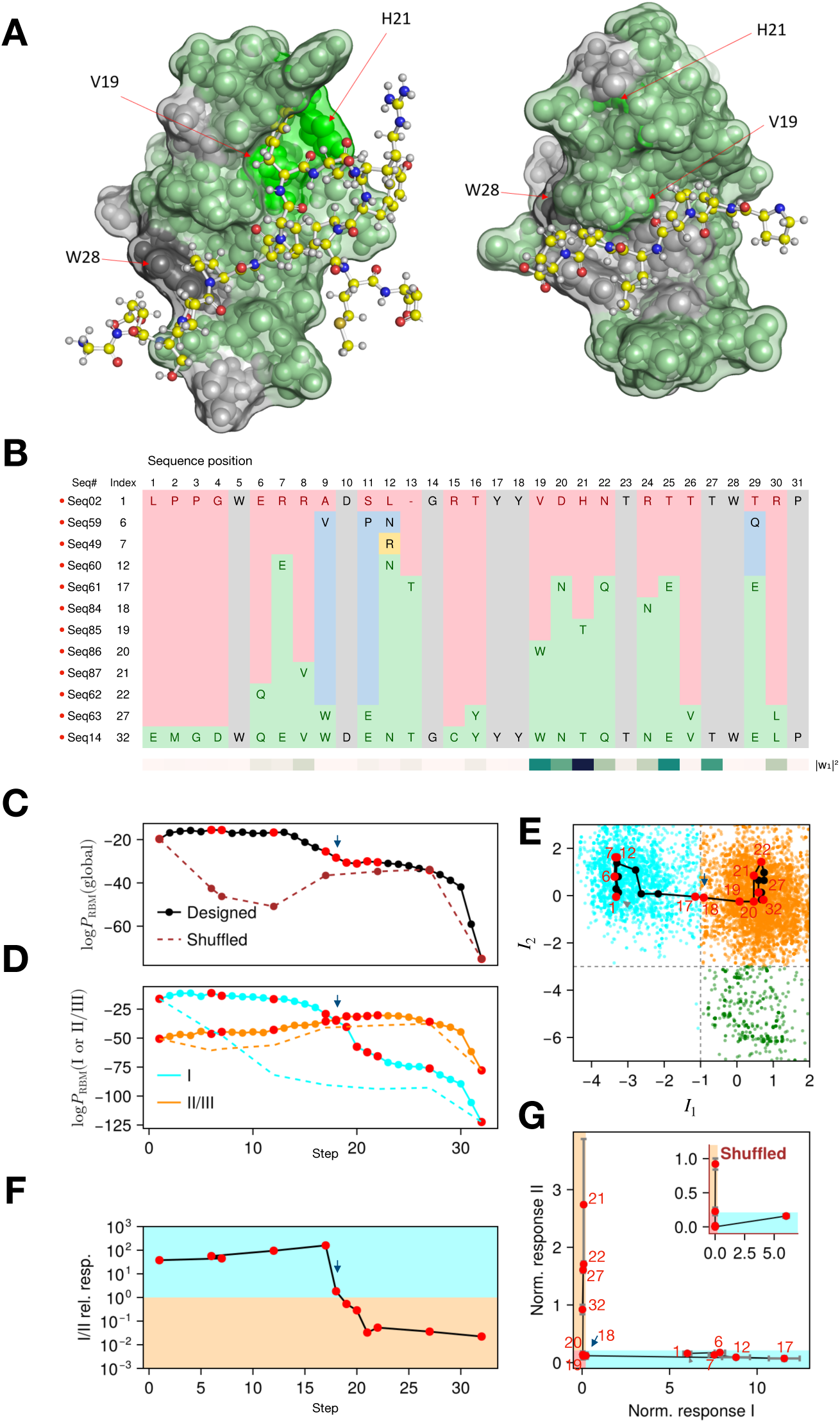
Path from Class I to Class II/III. **A.** Complexes of WW domain and cognate peptide of Classes I NP595793.1 (Pub3) (Left) and II 1YWI (Formin-binding protein) (Right); structures were predicted with Colabfold. **B.** Alignment of the tested sequences, showing the mutated amino acids. Same color code as in Figure 2. Sequence identity number (Seq#) and index (Index) along the path are specified. **C, D.** Global and class-specific RBM scores of the sequences in the path. **E.** Projection of the path one the plane of RBM latent units relevant for class identification. **F.** Ratio of experimentally measured responses to peptides of class II over Class I for intermediate sequences along the path. **G.** Normalized responses of the designed sequences against peptides of Classes I and II. The colored bands, defined by the experimental noise limit, indicates the non-binding zone.

To confirm that the order in which mutations are introduced along the designed path is dictated by epistatic interactions, and thus depends on the amino-acid background, we study “scrambled” paths, in which mutations are introduced in reverse order (Suppl. Table 6). This choice maximizes the sequence divergence between the background sequence (in which the mutation is predicted to be functional by the path sampling algorithm) and the one in which it is tested in the scrambled path. We observe that scrambled paths have global scores smaller than designed paths (Figure 3C), and do not contain any functional sequence (apart from the anchoring natural WWs), see inset of Figure 3H. Of notice, epistasis effects are correctly detected by Class I specific scores, but not by Class IV specific scores that assign a similar score to scrambled and designed paths. This observation is compatible with the absence of epistasis (Suppl. Figure 3) in the Class IV specific RBM model, probably due to the the scarcity of training data, but could also be related to a lower stability than Class I domains. To better understand the correlation between the two binding modes towards Class I and Class IV ligands, we study the specificity of sequences with adequate pairs of amino acids in position 24 & 25 for Class I specificity, namely RA or RT, or a mutated pair, NA, decreasing the binding affinity to Class I peptide, as well as long or short *β*_1_ − *β*_2_ loop. Fig. 3I, shows that that extending the *β*_1_ − *β*_2_ loop is not sufficient to achieve Class IV specificity, <u>e.g.</u> the sequence with RT in P24-25 and a long loop is characterized as being in Class I. Moreover, decreasing the affinity of Class I with the NA mutations and extending the loop is necessary to bind class IV ligands, underlining the presence of a long-range interaction between the two binding modes.

As shown in Suppl. Table 2,3,4, we have tested 3 additional paths interpolating between Classes I and IV. Remarkably, even on the path starting from the YAP1 wild-type, classified as non binding by our setup, all the designed sequences are functional (Supp. Table 2). All paths are found to follow similar trajectories in the two-dimensional plane, and show cross-reactivity in the region of the input space devoid of natural sequences, see Suppl. Figure 10: several sequences (Seqs. 52, 53, 54, 34, 35) even show mild cross-reactivity to Class II peptide. All paths pass through sequence ’WW150’ before going directly to ’WW10’ in 2 mutations, showing how constrained is the convergence towards Class IV reactivity.

We have also designed a direct path interpolating between the two classes (Suppl. Figure 10), along which transient mutations are forbidden. This path shows smaller RBM global score, especially for Class I sequences, in agreement with Section 3.2. Experimentally measured responses are very similar to the one found for indirect paths (red: no response; green: large response). A slight decrease in the binding response to peptide C1 is observed (Suppl. Table 4), but our experimental detection, focused on binding responses is not tailored to detect small differences in stability, see Section 3.2. In the region lacking natural sequences, the direct path goes through the same sequences (Seqs. 53, 35, 48) as the indirect ones, again underlying how constrained sequences are when approaching Class IV.

### 3.4 Paths from Class I to Class II/III

We now report results on paths interpolating between Classes I (wild type NP595793.1 Pub3) and II/III (Wild type 1YWI or Formin-binding protein1, also known as FNBP1, see Figure 4. This path critically differs from the Class I *→* IV case above as ligands of Classes I and II bind to the same pocket of the WW domain [12, 43, 11, 14] (Figure 4A).

Figure 4B gives the list of experimentally tested sequences obtained by subsampling the complete path given in Suppl. Fig 13. Note that the natural Class II/III domain at the end of the path (Seq.14) has two flanking sequences (2 amino acids in the N-terminus and 6 in the C-terminus), which are not covered by the RBM model. To study a path ending at Seq. 14 we thus added the same flanking regions to Seq. 2, leading to Seq. 2x with no loss in specificity, as well as to all the designed sequences along the path. Figure 4A shows the complexes formed by the WW domain and the two ligands; the majority of sites (*↑* 75%) in the molecule undergo mutations. Several residues are transiently mutated at the beginning of the path. Some of them (9-11-12) were already encountered on the path within Class I above, and increase the global scores (Figure 4C) with respect to the wild-type WW domain.

The global and specific scores relative to Classes I and II/III of the sequences are shown in Figure 4C,D. Notice that the natural WW domain of Class II/III at the extremity of the path has much lower global and specific scores than its Class I counterpart. This may be due to the fact that Class II/III domains are generally less stable than in Class I, as suggested by the contact map obtained from Class II/III specific RBM (Suppl. Figure 3) and by the lower conservation of residues in Class II/III domains, see sequence logo in Suppl. Figure 1 and [48] (Suppl. Figure 3. We predict a change in specificity after the fifth sampled sequence (Seq. 61) from the low-dimensional representation of Figure 4E and from the crossing of the specificity scores in Figure 4D.This switch requires mutations of all specificity-determining residues: V19W, H21T [12, 48, 43] and R24N, compatible with the weight vector associated to *I*_2_, and sequentially introduced from Seq.84 to Seq.86 along the path.

The normalized responses of the tested molecules are shown in Figure 4F,G. To better characterize the switch, we investigate the responses of the four designed sequences following Seq. 61, with index 17 in the path (crossing the specificity threshold in Figure 4E) to the three peptides (C1, C2, C3). No response to C2 peptide is found until the complete mutations of the three key residues above in Seq. 87 (with index 21), showing the largest C2 binding response, coinciding with the maximum of the Class II/III score. Previous rational design-based experiments for switching the specificity of Class I YAP (with peptide PPPPXP) WW to Class II/III FE65 (with peptide PPPPPP) have already pointed out that the two substitutions L19W and H21G were sufficient to perform specificity switching in the YAP1 background having Q24. As already mentioned, YAP1 is not responsive in our assay. Moreover, given the large fraction (6/15) of non responding wild-type sequences of class II, the response to the C2 peptide seems harder to detect. Lastly, we do not exclude that other non-tested ligands could be able to bind to such sequences, eventually in a less specific and more cross-reactive way.

We next experimentally analyze a set of ’scrambled’ sequences (Suppl. Table 6). These scrambled sequences have RBM specific and global scores smaller than the designed ones, and none of them is functional, see Figure 4G inset.

We report experimental results obtained for a second path switching specificity from classes I to II/III in Suppl. Table 5, see Suppl. Figure 10 for a low-dimensional visualization of the path. All tested sequences but one are responsive and bind peptides of Classes I or as predicted by the class-specific scores.

### 3.5 Comparison of model scores and binding measurements

The analysis of the mutational paths above has shown the ability of our computational model to generate new sequences behaving as WW domains with desired binding specificities. To further quantify this capability, we study the correlations between the global RBM scores and the experimentally assessed binding responses in Figure 5. The scores of the tested sequences, either natural or designed, have large values, biased towards the right tail of the histogram of natural WW sequences (Figure 5A). In particular, most designed sequences are, according to the global RBM model, as good as, or better than the majority of natural sequences. This result was expected as we set the effective temperature to the value 1*/*3 (lower than the standard value 1) to design sequences with high scores [34].

**Figure 5:**
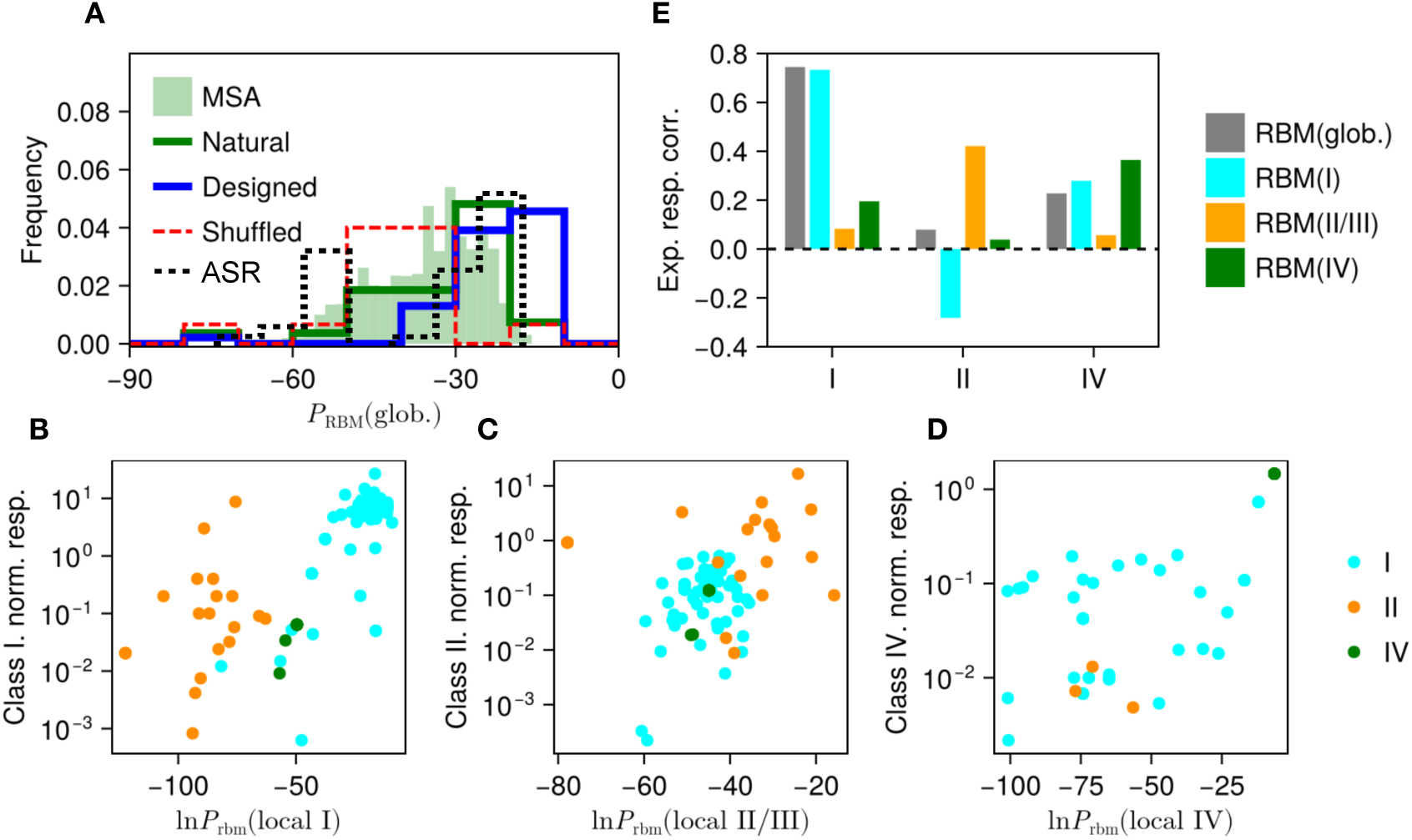
Model scores and experimental responses. **A.** Histogram of global RBM scores of probed sequences (Natural, Designed, and Shuffled). The histogram of global RBM scores of the full sequence data is also shown for comparison. **B, C, D.** Comparison between RBM class-specific scores and binding responses, for Class I (B), II/III (C), and IV (D) ligands. Sequences in each panel are colored according to the predicted classes, *i.e.* to the quadrants in the 2D projection plane defined by the specificity-related latent units. **E.** Pearson correlation coefficient between experimental responses to the three ligands (I, II, IV), and the RBM scores (specific and global).

Sequences along the paths obtained through reshuffling of the mutations in Figures 3 and 4 have, on the contrary, smaller global scores, in agreement with the experimentally observed lack of functionality. The scrambling of mutations, irrespectively of the amino-acid background, is expected to be detrimental in the presence of epistatic interactions.

Histograms of scores for the natural sequences associated to specific classes are given in Suppl. Figure 1 and confirm the RBM scores of Class I natural sequences have typically larger values than the ones of Classes II/III and IV, which could be indicative of stronger structural stability.

In Figure 5B,C,D, we compare, for each class, the scores assigned by the specific RBM models to the experimental binding measures of the corresponding class ligand. We observe positive correlations, see Figure 5E for the three models, and less so for the global score. Class I is an exception, as the global score also shows a strong correlation; this may be due to the fact that class I WW have the larger scores as shown in Suppl. Figure 1 and several contacts are kept by conserved amino acids in Class I domains [48], see logo representations and contact map for Class I in Suppl. Figure 3.

### 3.6 Comparison with Ancestral Sequence Reconstruction on WW domains

Lastly, we compare the mutational paths designed with our method between wild-type domains with the pathways linking these domains on the phylogenetic tree derived from the WW domains present in the PFAM seed. To do so, we reconstruct the tree connecting all these domains and perform Ancestral Sequence Reconstruction (ASR) on the intermediate nodes. This tree topology is inferred directly from the WW domains only [52] and not from the proteins they belong to. Multiple pathways connect the domain subtypes on the tree (see Suppl. Figure 16). As illustrated in figure 6A and with the collapsed tree topology in figure 6A, we observe that all type I and IV domains are connected through a narrow path, which crosses the empty quadrant in the 2D-latent space defined by the two specificity-related RBM hidden units. This result is similar to what is obtained with our RBM-based method.

**Figure 6:**
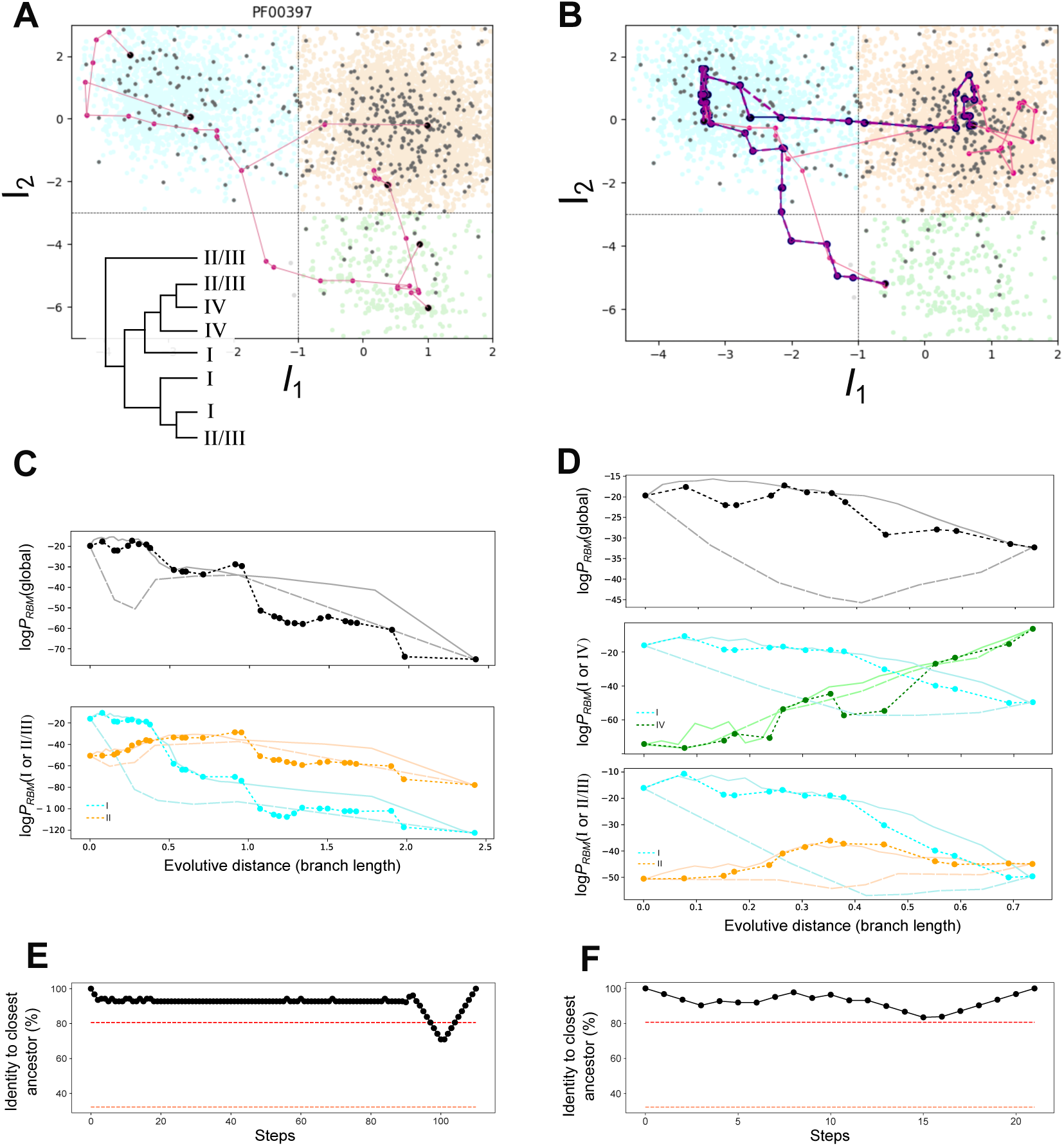
Pathways derived from ancestral sequence reconstruction (ASR) on the WW domains in PFAM’s seed. **A.** 2D projections of selected paths linking domains of different types (I, II/III and IV) on the two specificity-related RBM hidden units, and simplified representation of the seed tree topology. Black dots: modern sequences; Purple dots: average ancestral sequences sampled from the posterior distribution. **B.** 2D projections of paths linking WT domains presented in Figures 4 and 3 along the tree. Black: mutational paths designed with the RBM; Purple: paths obtained with ASR; Dotted purple: ASR path obtained when adding to the WW tree the RBM intermediates. Blue and purple paths are largely overlapped with one another. **C., D.** Scores of ancestors obtained with ASR (dotted lines) and with the global (top) and specific (middle panels) RBM models. Scores obtained along RBM-sampled paths and scrambled sequences are shown as plain and dashed lines for comparison. **C.** Path I to II/III. **D.** Path I to IV. **E., F.** Identity between intermediate sequences along the RBM path and the closest ancestors reconstructed by ASR; median and maximum identity between seed WW domains are shown as dashed red line for reference. **E.** Path I to II. **F.** Path I to IV.

While ASR paths have in median a large number = 113 of mutations (Suppl. Figure 17, middle bottom panel), RBM paths were constrained to be much shorter, with 14 to 25 mutations, a length shared by few ASR paths only, see Suppl. Figure 10A, B, C. This observation could explain why RBM-designed paths cross a region devoid of contemporary sequences. Remarkably, while ASR paths are not constrained in length, the pathways connecting type I and type IV domains also follow the same narrow path. Notice that the phylogenetic tree inferred from WW domain sequences only directly connects domains of type I to those of type IV, different results are found with trees obtained from families of homologous proteins containing WW domains as a subpart of their sequences. In the latter case, paths going from I to IV pass through region II/III (Suppl. Figure 18) In particular, the phylogeny of the protein NP001351424 (peptidyl-prolyl cis-trans isomerase from *Mus musculus*), which contains the WT class IV domain used as anchor for path sampling, shows intermediate ancestors between class I and II/III homologs belonging to the class II/III (Suppl. Figure 18D).

The funnel connecting directly WW domains of classes I and IV is therefore an interesting feature of WW-domain reconstructed trees, not locked to the evolution of the rest of the protein, and possibly better reflecting the selective pressure associated with binding specificity. The differences between the topologies of the trees found from the WW domains and from the proteins containing them are compatible with previous observations [52]; in particular, domains with different specificity types in the same protein may be less related to domains of the same type on different proteins.

Figures 6B shows that the ASR sequences along the pathways (in purple), connecting the wild-type sequences anchoring the RBM paths are strikingly similar with the RBM intermediates. Their residue identity exceed 90%, as evidenced in Figure 6E and F. Conversely, the median identity between randomly selected modern domains in the seed is 33%. The scores of the ancestral sequences along the phylogenetic pathways assigned by the RBM are larger than the ones of scrambled paths, and slightly but consistently lower than the ones of RBM-designed sequences (Figures 6C and D), see Suppl. Fig 6 for comparison of the scores of the most ancestral nodes along the paths with the consensus sequence of the wild-type WWs and the distribution of natural scores. This result is expected as ASR reconstruction does not take directly into account epistasis, while RBM may account for multiple-residue correlations.

Incorporating the RBM-designed sequences into the phylogenetic tree further corroborates their similarity with the ancestral sequences. These RBM sequences collapse on the trajectory connecting the wild-type sequences (Figure 6B, with the dotted purple ASR paths overlapped on the blue synthetic paths).

## 4 Discussion

In this work, we propose and test a computational method to design mutational paths interpolating between WW domains binding classes of proline-rich peptides [18, 53]. Our approach relies on a model of the fitness landscape of WW domains learned by Restricted Boltzmann Machine (RBM) from homologous sequence data and on Monte-Carlo sampling [30, 44] of functional paths [26] that connect extant WW domains across sequence space. Using a high-throughput in-vitro approach, we test the binding responses of the designed sequences toward three peptides representative of the three main WW specificity classes. Our study provides experimental validation of multiple designed paths both within the same WW specificity class (binding the same peptide) and across different specificity classes, capable of switching specificity through accumulation of mutations. These experiments, in turn, give insights on the main requirements for the change of specificity in a WW protein domain. We finally compare the RBM-designed paths with pathways obtained by ancestral sequence reconstruction (ASR) on a phylogenetic tree built on the WW domains of the PFAM seed.

Navigability in the fitness landscape was first studied in evolutionary biology using theoretical models [18, 54, 55, 56]. In recent years, the availability of massive experimental data began to allow for partial fitness landscape reconstructions. The approach introduced here for path design is, in a way, a synthesis between these two approaches. Unlike theoretical frameworks such as the ’house of cards’ [57] or the NK [54] models, our fitness landscape is directly inferred from sequence data and is therefore, at some level of approximation, representative of the complexity of a real protein domain [43, 30, 58]. Moreover, our approach borrows from statistical physics some conceptual and methodological ingredients, such as Monte-Carlo sampling of paths [54]; in addition, the bipartite nature of the RBM network makes it possible to use mean-field approximations to theoretically characterize the topology and accessibility of the fitness landscape [54, 23], e.g. the mean trajectory of paths and their entropy [44, 59]. Computational approaches enormously reduce the huge number of a priori paths to be experimentally tested, and could speed up the rational design of protein selectivity, attempted since the 1990s in the field of drug discovery, in which few mutations targeting key specificity residues were probed, often ignoring the presence of complex epistatic effects between residues [14, 60, 61, 62, 63].

Overall the designed paths are rich in functional sequences: only 4 out of 58 tested sequences were non responsive to any of the three tested ligands, while all the other sequences responded to at least one of the peptide representative of the specificity class corresponding to the natural WW domain at the extremities of the path. This result proves the general validity of combining generative models for the fitness landscapes inferred from sequence data and Monte-Carlo path sampling to propose viable paths. It extends the recent success in protein design with computational methods [64] to the design of full mutational paths. Our RBM model is capable of capturing important epistatic effects constraining the order in which mutations can be introduced along the path. As shown by scrambling the mutational order along the path, sequences in which mutations take place independently from the background content are not functional, as they do not take into account epistasis. We stress that the modeling of epistatic interactions with the RBM is global over all WW domains, and not attached to any specificity class. It is not possible to learn accurate landscapes for each single specificity as only *↑*50 out of 17,000 available WW sequences in databases have been experimentally characterized in terms of binding specificity [11].

Being shallow, the RBM parameters of the model are interpretable, and unveil both key specificity determining residues, similarly to methods based on principal component analysis ([65, 66]) and epistatic interactions as in the Direct Coupling Analysis [67, 68]. Moreover, the tested specificities along the paths are in good agreement with the approximate class-specific RBM scores we have introduced and specificity switches take place through changes in specificity determining residues and amino acids identified by the RBM weights attached to the two hidden units identified as most informative about specificity classes [43]. The identity of the binding class is therefore correctly predicted by the clustering of the sequences generated along the path according to these hidden units, as well as by the class-specific RBM scores.

Our designed paths allow for the presence of ”transient” substitutions with amino acids different from the ones in the anchoring wild-type WW sequences [69], differently from [21]. In all the paths we tested, we observe such mutations to transient amino acids on variable residues away from the main specificity determinant. These mutations are important: they transiently improve the global RBM score in the initial part of the path, by possibly increasing the stability of the molecules with respect to the class I wild-type WW domain, while preparing the background for the specificity change. However, class switching typically involves mutating the specificity determining residues to the amino acids found in the final wild-type WW domain. The above findings are in full agreement with the stability-specificity tradeoff previously introduced in evolutionary path characterization [56, 70, 71] and in the specific characterization of Pin WW variants [72].

As the binding pockets of peptides is shared for Classes I and II, and differs for Class IV, it is expected that specificity-switching paths from Class I to II and from Class I to IV be qualitatively different. Our study shows that Class I to Class II switch is more abrupt with a binding loss towards the particular peptides we have experimentally probed taking place roughly half-way along the path. Binding is restored once three specificity-determining residues are mutated. Conversely, along class I to IV paths, specificity can switch (steps 20-21), with no complete loss of affinity with the peptide of class I. To assess if the two binding regions are really independent or not, we have tested some additional specific mutations along the class I-IV mutational paths. In our attempts to engineer cross-reactivity, we have observed that it is important to substantially lower affinity to class I peptide to acquire class IV specificity, in agreement with previous studies [72].

Mutational paths connecting classes I and IV cross a region deprived of natural sequences in the low-dimensional projection defined by the two specificity-related RBM hidden units, see Figure 3F [30]. Synthetic WW domains along the I-IV paths go through a bottleneck in the space of sequences: several designed and tested paths go through the same intermediates, and are characterized by cross-specificity to peptides of Classes I, II and IV. Ancestral promiscuity and switch of ligands have been proposed for several enzymatic families based on ASR [17, 73, 29]. Promiscuous WW domains could be related to ancestral states, which could have specialized during evolution in favor of more stringent ligand selectivity. In Humans, WW domains exist also in tandem and multi-domains, and gene duplication is a plausible mechanism for such a specialization [49]. Reconstruction of ancestral sequences (ASR) from domains contained in PFAM’s seed supports this hypothesis, unveiling a similar narrow pathways connecting type I and IV domains, with ancestral nodes showing a large residue identity with the RBM sequences along the path. We stress that phylogenetic reconstruction made at the protein level –and not at the domain level– for WW-domain-containing protein families rather shows transitions between classes I and II/III and classes II/III and IV. More work is thus needed to assess the evolutionary significance of the putative ancestral sequences along the I-IV path found in the WW-domain tree.

ASR and RBM-path sampling are complementary tools to explore hypothetical paths between distantly related modern sequences, both based on hypothesizing evolution as a parsimonious path in a fitness landscape. This analogy between the fitness landscape inferred from sequence data and the phylogenetic paths has already been used in protein engineering as a method to infer novel proteins with enhanced properties such as affinity, stability, and solubility [74, 75] and to predict mutational effects in [76], where the distance between sequences on a Neighbor-Joining tree serves as a proxy to assess the likelihood and, consequently, the fitness of novel mutations in a protein. In the context of WW domains, both ASR and RBM models propose the existence of a narrow path of domain evolution connecting type I and type IV domains.r ASR internal nodes have scores comparable even if slightly worse than RBM sampled sequences along the path. However, ASR offers much lower resolution in the sequence space compared to the RBM, as the computational complexity in reconstructing a tree makes impossible to use all sequences available in the databases; it does not provide a true quantification of the likelihood of the intermediate sequences and does not model epistatic interactions between residues. The similarity of ASR and RBM paths proves that the sampling of paths in the fitness landscape with the RBM could help to explore likely intermediate sequences between groups of distantly related proteins, a task for which classical ASR is not well suited, and which could prove useful both for protein engineering and phylogenetic studies.

Our work could be extended in several directions. First, we plan to extend our approach to sample paths attached not at their two extremities, but anchored to one wild-type WW domain only. Imposing that the affinity to a ligand different from the initial one increases along the path would make the setting more closely matches a natural evolutionary path. Additionally, it would be worth investigating longer and more complex proteins, for which we expect the disentanglement of specificity-switching latent units [77] to be more difficult. However, while relating the latent units to specificity classes was convenient in the present study, it was never used to generate the paths themselves. As an illustration, we have recently used the same approach to sample viral trajectories, more precisely, variants of the Receptor Binding Domain of the SARS-CoV-2 spike protein capable of escaping antibody recognition[59]

Second, the in-vitro assay could be complemented in several ways. Our current assay does not allow for cooperativity between WW domains, which may be important in vivo. This could explain why some natural WW domains tested alone, such as YAP, do not display affinity to their cognate ligands. It has been previously shown that multi tandem WW domain confer higher ligand affinity [49]. In addition, our current experimental setup is limited to a small number of peptides. It is possible that the four non-responsive sequences we have sampled on the path connecting Class I and Class II/III wild-type WW domain may bind to other non-tested peptides representative of the same classes, possibly in a less specific and more cross-reactive way. To better investigate the above points we plan to extend the experimental binding measures to in vivo binding assays, and to measure binding against a library of peptides through competitive growth in yeast cells [12, 78]. Besides, a more exhaustive experimental characterization of designed sequences along direct versus non-direct paths, in particular along the class Class I to Class II/III specificity switch, would be interesting to investigate the gain in exploiting the full amino-acid space in the transition path. Last, while we have here used experiments as a way to validate the paths proposed by the computational method, it would be interesting to iteratively improve our estimate of the fitness landscape inference by integrating the new experimentally labeled data using active learning methods [79, 80, 81, 64, 82].

## 5 Materials and Methods

### 5.1 Restricted Boltzmann Machines

We train our unsupervised model on a data-set of homologous proteins, presented as a list of aligned sequences {**v**^1^, **v**^2^,…, **v***^k^*,…, **v***^B^*} (*B* is the total number of sequences in the data set), where each entry is an array 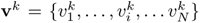 in which each one of the *N* entries may be in one of the possible 21 states (20 amino acids + the gap site). As a model, we use Restricted Boltzmann Machines [83], a neural network consisting of a visible layer **v** representing the data and an hidden layer of *M* neurons 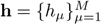. RBMs define a joint probability distribution over the visible and hidden layer as

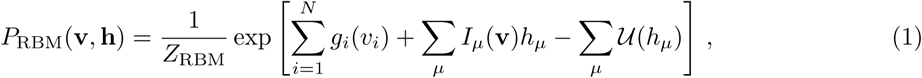

where *Z*_RBM_ is the normalisation constant, the input to an hidden unit is the projection of the sequence on the weigth matrix *w_i,µ_*: *I_µ_*(**v**)= Σ*_i_ w_i,µ_*(*v_i_*) and *U_µ_* are potential energy functions over the real-valued hidden unit activations *h_µ_*. We choose *U_µ_* to be double Rectified Linear Unit (dReLU) potentials of the form

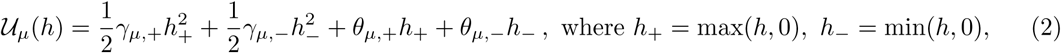

where we have defined the hyper-parameters *γ_µ,±_*, *θ_µ,±_*. Marginalising over the hidden units we obtain the probability distribution over the sequence space

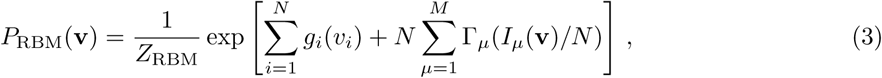

where 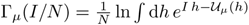. We use Persistent Contrastive Divergence [84] to train the model over the multi-sequence alignment of the protein family of interest in order to maximise the likelihood. This training algorithm has been shown to be sufficiently robust under cautious regularization [85]. The code and the data used to train our RBMs for WW domains can be found in [86], while the hyper-parameters for the training can be found in the Supplementary Materials of [30].

### 5.2 Global and specific RBM models

We learned two classes of RBM models, depending on the train data. A global RBM model used to design the paths was learned on all the sequence data, while three class-specific RBM models, for binding classes of type I, II/III, or IV were learned on sub-set of sequences (containing 8304, 8292, 637 sequences respectively). To define the sub-sets we clusterized the sequences according to the value of their inputs *I_µ_*(**v**) on the two hidden units *µ* = 1, *µ* = 2 chosen as the most informative about the specificity classes, shown in Figs. 2, 3, 4. The specificity threshold indicated in the figure (*I*_2_ < −1; *I*_2_ > −1, *I*_1_ < −3; *I*_1_ > −3) have been informed in [30] by some previous experimental tests [12, 11] (see Figure 3 in [43] and Figure 2a in [30]) on the WW specificity classes. Compared with other prediction methods such as *k*-means and variational auto-encoders, the RBM classification achieves a strong prediction accuracy (94.7 %) compared to experimentally assessed domains as reported in Figure 2E and Suppl. Figure 7. The performance of the different methods are presented in Suppl. Figures 8 and 9. The clustering according to the RBM latent space results in three class-specific RBM models (one for class I, one for class II/III and one for class IV) which can be used as predictor of the binding affinity of a natural or artificial sequence.

### 5.3 Path sampling algorithm

We define the probability a path of *T* steps, *V* = {**v**_1_, **v**_2_ …, **v***_T-_*_1_}, connecting two sequences **v**_start_ and **v**_end_, as follows:

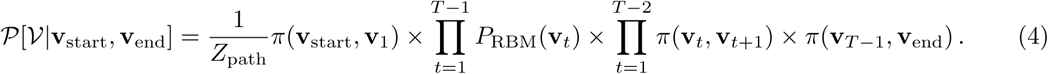

Here *π* is an interaction terms that is used to minimised the number of mutations at each step. Our goal is to sample paths that do at most one mutation at each time step. In practice we set *π*(**v**, **v**′)= 1 if the two sequences are equal, *π*(**v**, **v**′) = *e*^−⋀^ (with ⋀ *>* 0) if the two sequences differ by one mutation and *π*(**v**, **v**′) = 0 otherwise. In [30] we propose a Monte-Carlo (coupled with simulated annealing) procedure to sample paths from *P*(*V*)*^β^* at a given inverse temperature *β* = 1*/T*. Other choices of the transition term *π* are discussed in the following subsection.

Here we stress that our path sampling approach is independent of the restricted Boltzmann machine used to train the model and one can use in principle different probability distribution *P*(**v**). An alternative choice would be to define our path sampling algorithm over the space of nucleotide sequences that encode for the specific protein. Each sequence would be scored according to the RBM model after translation into the corresponding amino acids sequence according to genetic code. In this way one can introduce codon biases in the sampling approach. We applied this approach to the case of WW domain and we obtained similar results to the sampling procedure carried out without codon bias.

### 5.4 Ancestral protein reconstruction on seed domains

Phylogenetic tree inference and Ancestral Sequence Reconstruction (ASR) were conducted to obtain an unrooted tree topology and ancestral node sequences for all 408 isolated WW domain in PF00397. We used IQ-Tree [87] and an evolutionary model defined by the amino-acid mutation matrix LG [88] and Gamma parameters discretized in four classes [89] (LG+G4, identified as the best model to describe domain evolution, see Supplementary Information, Section D.2). Gaps were reconstructed from the sequence alignment converted into a binary presence/absence matrix from binary sequences, using the GTR2 model [90]. We calculated inputs to the hidden units of the RBM for modern WW domains, as well as their mean values for ancestral sequences based on the posterior probabilities of ancestral residues. Scripts to perform the ASR are available on the project’s Github.

We also performed phylogenetic reconstruction from the 24 seed domains within or near cluster associated with type IV domains, as well as the NP00131424 protein used as a type IV anchor for the RBM path sampling, as described in Supplementary Information, Section D.2.

### 5.5 Experimental Materials

Cell free expression system (PURExpress ^®^ In Vitro Protein Synthesis Kit, New England Biolabs, E6800), SNAP labeling Dye (SNAP-Surface^®^ Alexa Fluor^®^ 647, New England Biolabs, S9136), Anti-SNAP antibody (SNAP-tag Polyclonal Antibody, Thermo Fisher Scientific, CAB4255), Streptavidin Magnetic beads (Dynabeads™ M-280 Streptavidin, Thermo Fisher Scientific, 11205D), Protein A magnetic beads (Pierce™ Protein A Magnetic Beads, Thermo Fisher Scientific, 88845), oligonucleotide linkers and eBlocks™ Gene Fragment provider (Integrated DNA Technologies), synthetic peptides provider (GenScript Biotech), Gibson assembly (Gibson Assembly^®^ Master Mix, New England Biolabs, E2611), Polymerase kit (Q5^®^ High-Fidelity DNA Polymerase, New England Biolabs, M0491) DNA purification beads kit (SPRIselect^®^, Beckman Coulter, B23318) and phosphate-buffered saline buffer (DPBS 10X, Thermo Fisher Scientific, 14200075), DTT (DL-Dithiothreitol solution, Sigma-Aldrich, E2611), Tris-HCl (UltraPure™ 1 M Tris-HCI Buffer, pH 7.5, Invitrogen, 15567-027), Tween-20 (TWEEN^®^ 20, Sigma-Aldrich, P2287), EDTA (0.5M EDTA pH 8.0, Invitrogen, AM9260G), Binding buffer (Tris-HCl 50 mM pH 7.5, EDTA 10 mM, DTT1 mM and 0.5 % Tween 20, prepared in-house), Wash buffer (1X DPBS with 0.5% Tween-20).

### 5.6 Experimental protocol

Our experimental setup is sketched in Figure 1D. Double-stranded DNA gene fragments encoding the WW domain sequences were synthesized and inserted by Gibson assembly into a plasmid containing a T7 promoter and a ribosome binding site (RBS) upstream of the cloning site, these elements being essential for initiation of the expression process. A sequence encoding a linker and a SNAP-tag has been fused downstream of the WW domain sequence and a stop codon has been placed at the end of the SNAP-tag sequence followed by a T7 terminator sequence. The SNAPtag is a protein capable of reacting irreversibly with the benzylguanine (BG) group. This can be used to label the protein fluorescently, using dyes coupled to BG.

After insertion of the WW domain genes, a polymerase chain reaction (PCR) was conducted to amplify the entire construct, producing a linear product encompassing the complete sequence between the promoter and the terminator. The resulting PCR product was then utilized for expression in a cell-free transcription and translation system, with the fluorescent substrate.

Concurrently with the expression of the WW protein, magnetic beads coated with the target peptides, which are presumed to bind with the WW domains, are prepared. These comprise a metallic bead displaying a multitude of copies of the peptide of interest on its surface, as well as a fluorescent dye for the identification of the beads. At the same time the protein A-coated magnetic beads are incubated with a polyclonal anti-SNAPtag antibody for the purpose of assessing the efficacy of the expression process.

Following the expression of the fusion proteins and labelling with the fluorescent dye, a mixture of color-coded beads coated with different peptides and anti-SNAPtag antibody was added to the expression mixture. Upon binding of the WW domain to the peptide, the fluorescent dye attached to the SNAP tag will report the amount of protein bound to the bead. The beads were then washed and passed through a flow cytometer, where the average dye fluorescence intensity on the bead surfaces was measured. This indicated the amount of WW domain fusion proteins bound to each peptide-coated bead and the expression assessment beads, which was then normalized for expression. Each WW domain protein was evaluated against three peptides, each representing a distinct class of WW domain. Further details are available in Supplementary Information, Section 5.6.

#### Fusion Protein assembly, expression and labeling

eBlocks™ gene fragments for genes comprised in the different pathways were designed, ordered, and cloned into the SNAP fusion plasmid using Gibson Assembly® Master Mix. Using Q5® High-Fidelity DNA Polymerase, PCR reactions were performed to amplify linear products containing all the necessary parts for the expression of the fusion proteins. PCR products were purified using SPRIselect® beads. Expression of the fusion proteins were performed in a PURExpress® in vitro protein synthesis reactions with the purified linear PCR products at 15 nM and SNAP-Surface® Alexa Fluor® 647 at 5 µM final concentration. The cell-free *in-vitro* transcription and translation (IVTT) reactions were incubated for 5 minutes at 37°C followed by 4 hours of incubation at 30°C.

#### Expression control beads preparation

Expression control beads were prepared using 5 µl of protein A magnetic beads (50 ug) from Thermo Fisher Scientific, washed twice with 1X DPBS from Thermo Fisher Scientific with 0.5% Tween-20 from Sigma-Aldrich, resuspended in 50 µl of 100 ng/µl anti-SNAP polyclonal antibody diluted in 1X DPBS from Thermo Fisher Scientific with 0.5% Tween-20 from Sigma-Aldrich, incubated for 30 min at 37°C with shaking, then washed three times with binding buffer.

#### Streptavidin beads preparation with target peptides

beads were prepared using 5 µl of streptavidin magnetic beads from Thermo Fisher Scientific, washed twice with 1X DPBS from Thermo Fisher Scientific with 0.5% Tween-20 from Sigma-Aldrich, then resuspended in 50 µl of 1uM DNA linkers from Integrated DNA Technologies diluted in 1X DPBS from Thermo Fisher Scientific with 0.5% Tween-20 from Sigma-Aldrich. The beads were incubated with the DNA linkers for 30 minutes at 37°C with shaking at 600 RPM in ThermoMixer® C from Eppendorf, then washed twice with wash buffer and resuspended in 20 ng/µl of target azide peptide from GenScript Biotech diluted in wash buffer, then incubated overnight at room temperature with rotation. After washing the beads twice with binding buffer, the beads were resuspended in 50 µl of binding buffer, after which beads with different targets and identification fluorescent dyes were mixed.

#### Binding

10 µl of labeled proteins were mixed with 4 µl of beads mixture then incubated for 1 hour at 25°C with shaking at 600 RPM in ThermoMixer® C from Eppendorf. After binding, the beads were washed twice with wash buffer and resuspended in 200 µl of wash buffer. Finally, the fluorescence intensity on different bead species was measured Guava® easyCyte flow cytometer (Millipore).

#### Experimental Design

Each WW domain was tested against three peptides, each one corresponding to a different class of WW domains. The three peptides are given in Table 1. IVTT No expression control and no binding controls (peptides with different sequences; IAKLDEESILKQ and PALWKSKGAD) was performed for each experiment to confirm the specificity of WW domains binding to the target peptides. All binding experiments, including target and control peptide measurements, were conducted in triplicate to ensure reproducibility and statistical robustness.

## Supporting information

Supplementary Informations

## 6 Data and Code Availability

All codes will be accessibles in https://github.com/orgs/CoccoMonassonLab/ and https://github.com/cossio/TransitionPaths2025.jl

## 7 Acknowledgments

This work was supported by the grants ANR-19 Decrypted CE300021-01, ANR-24 CE15-Methaflu and, ANR-23 CE45-0034 ProDiGen.

