## Supplementary Informations for "Design and experimental characterization of specificity-switching mutational paths of WW domains"

**A. Supplementary Figures**

**B. Tested sequences**

**C. Experimental protocol**

**D. Ancestral domain reconstruction on families of homologous proteins**

#### A Supplementary Figures

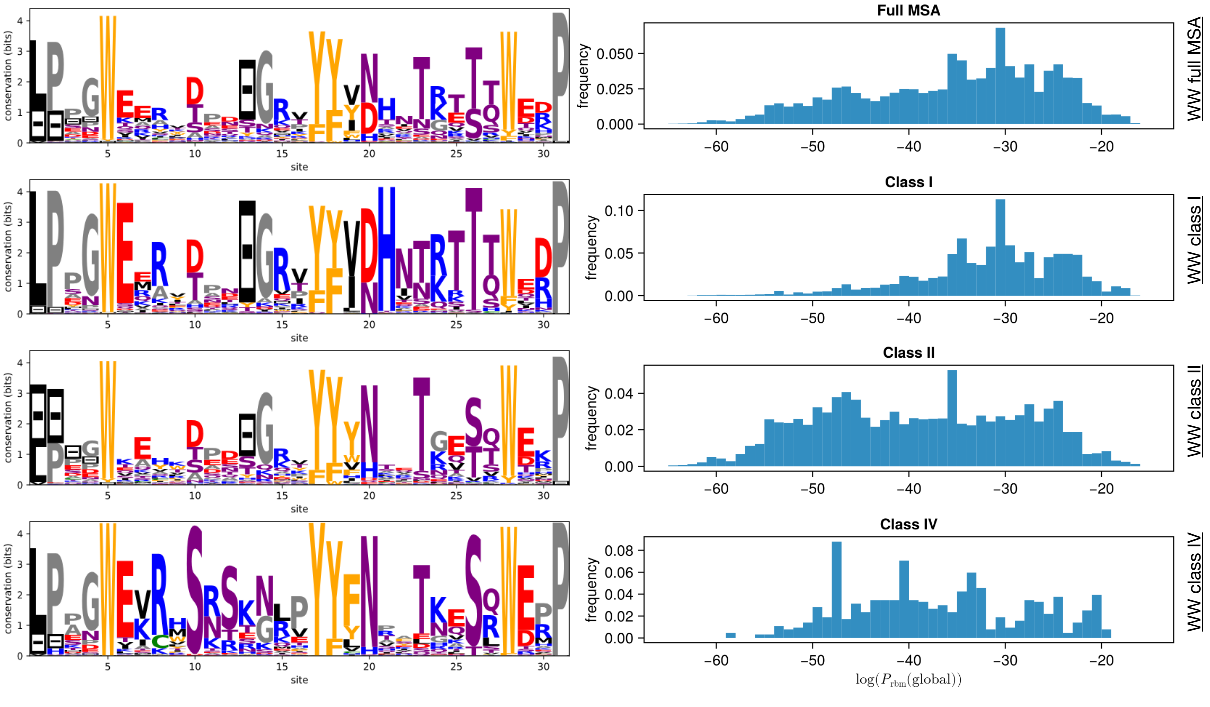


Suppl. Figure 1: Left: Sequence Logos of the Full MSA and the class specific MSAs, Right: Histograms of the RBM global scores for the whole MSA and each sub-sets. Class 1 has strongest conservation in particular on residues in contact. *E.g*, contacts 20-21, 21-26, and 6-21.

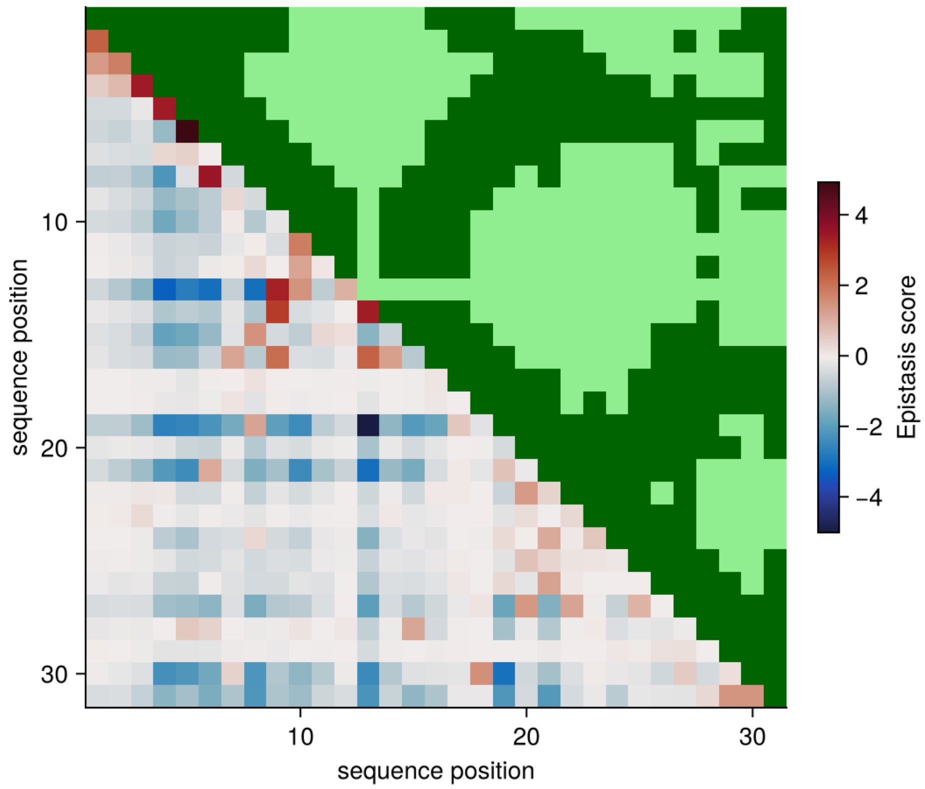


Suppl. Figure 2: Epistatic scores computed from the RBM weights [43], the pairs of residues with the highest epistasis scores are 6-8, 9-13, 9-14, 13-14, 8-16, and 9-16. The upper triangular part shows the real contacts (distance $<$ 8A) in the PDB 2LTW.

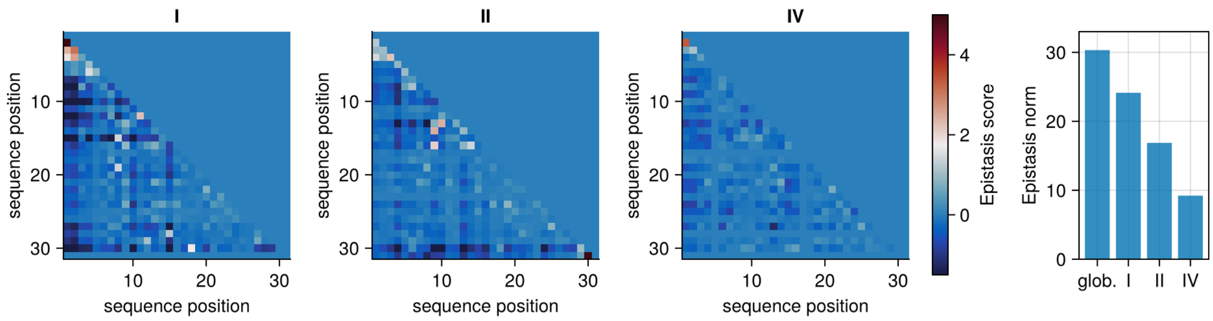


Suppl. Figure 3: Contact maps obtained from the three class-specific RBMs: 'I' trained on natural sequences falling on the specificity quadrant I, 'II' trained on natural sequences falling on the specificity quadrant II/III, and 'IV' trained on natural sequences falling on the specificity quadrant IV. Contact maps are computed as in Figure 2. The last panel on the right compares the norms of the contact maps (excluding diagonal entries) of the global and class-specific RBMs.

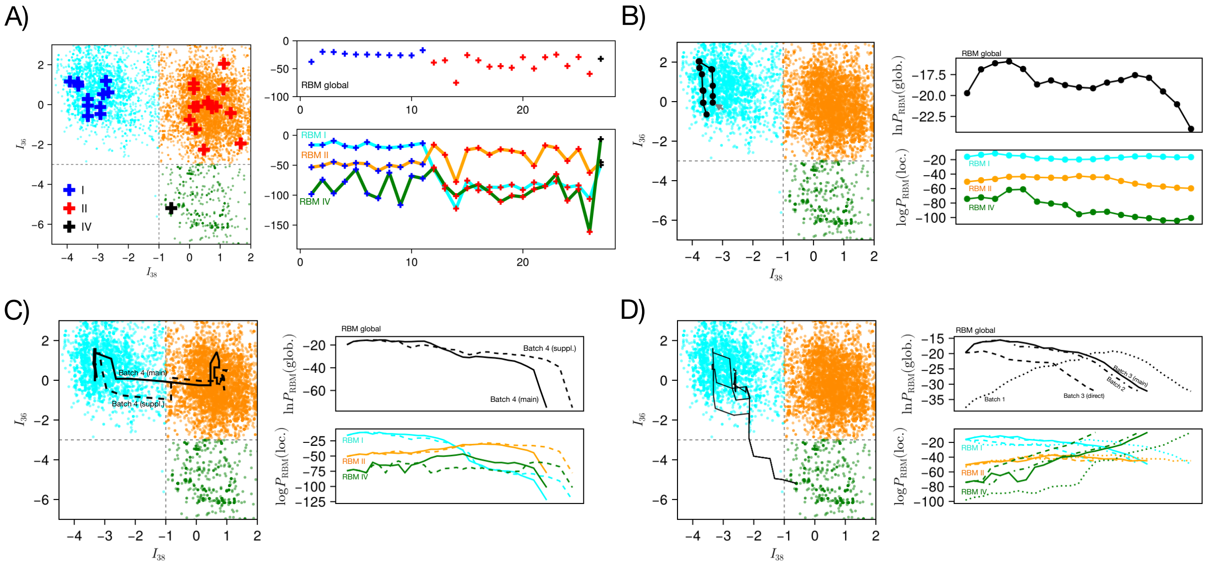


Suppl. Figure 4: **A)** Projections of the tested wild-types along the two specificity RBM weights (colors indicate the specificity label, known from literature [43]). Right: scores to the sequences according to the RBM global model (top) and type I,II,IV specific RBMs. **B)** same as A) for the I → I path (Batch 2, in the main) **C)** same as A) for the two paths from I → II/III specificity (Batch 4) **D)** for the 4 paths from I→ IV specificity (Batch 1, starting from the non responsive YAP1, Batch 2 , Batch 3, including the path in the main and the direct path, which is shorter and has a smaller global score at the beginning)

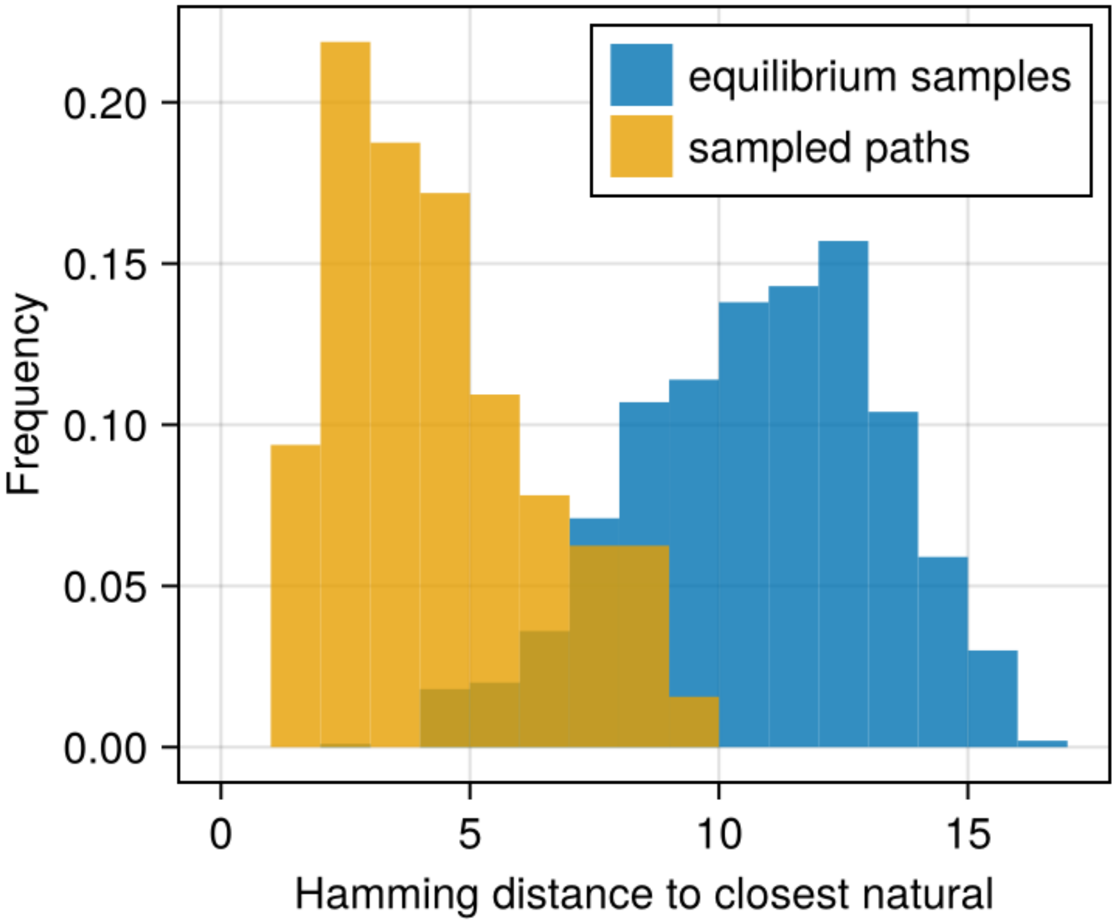


Suppl. Figure 5: Histogram of Hamming distances to closest natural sequences of equilibrium samples from the RBM (in blue). We also show the histogram of distances for all the intermediate designed sequences in the transition paths, to the closest natural sequence excluding the wild-types at the extremeties of the path (in yellow).

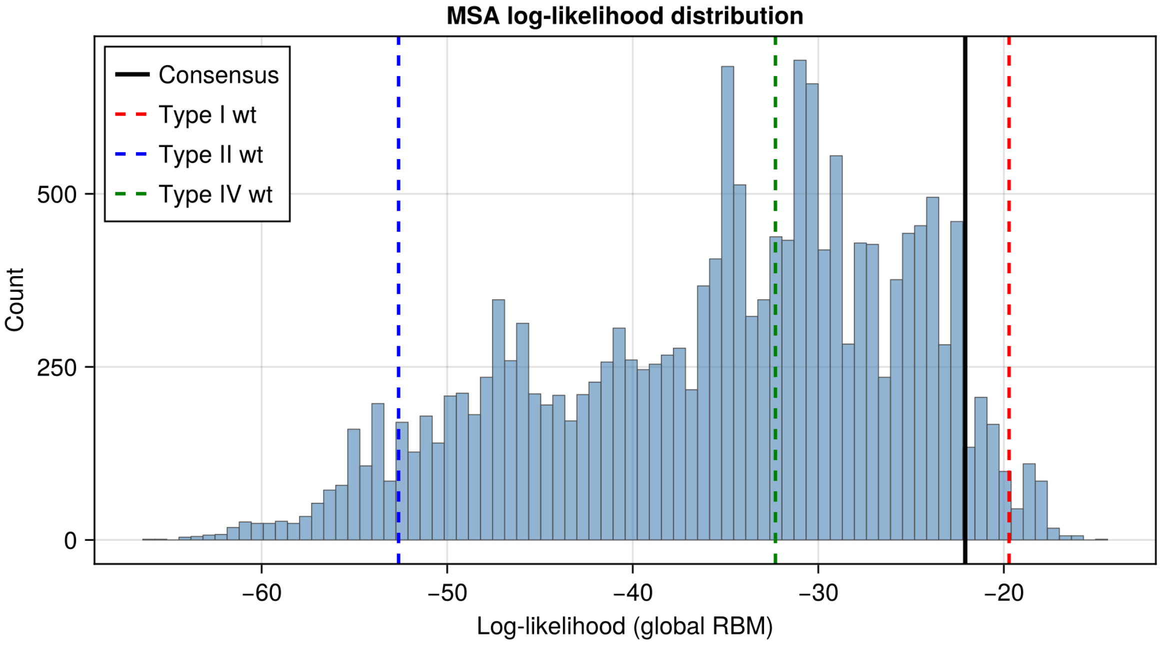


Suppl. Figure 6: RBM scores of the consensus sequence, of the wild-type extant sequences along the paths, and of the most ancestral ASR sequence along the ASR paths compared to the distribution of scores of natural sequences used to train the RBM model. The most ancestral nodes in the paths have 81% (paths between type I and IV domains) and 54%(paths between type I and II/III domains) similarity to the consensus, and an RBM score of -21, or -58, respectively.

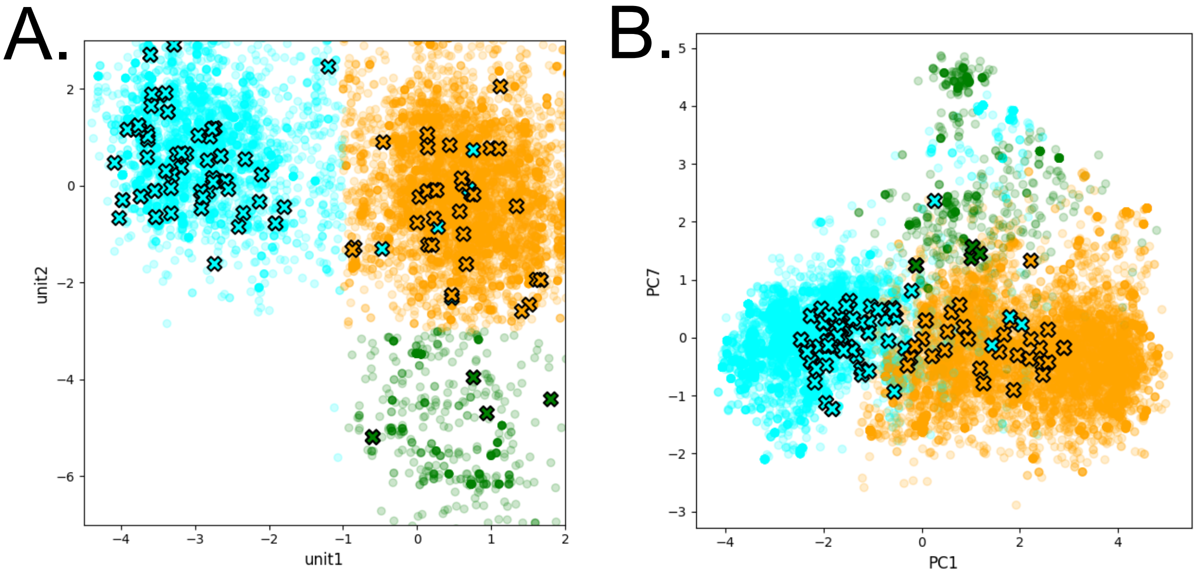


Suppl. Figure 7: Experimentally assessed sequences against RBM classification. Type I domains are in cyan, type II/III in orange and type IV as green. RBM-classified natural domains are represented as dots, and experimentally assessed sequences as cross [12, 11]. A. Projection against two hidden units inputs of the RBM. B. Projection against the principal component 1s and 7 of the one-hot encoded natural sequences.

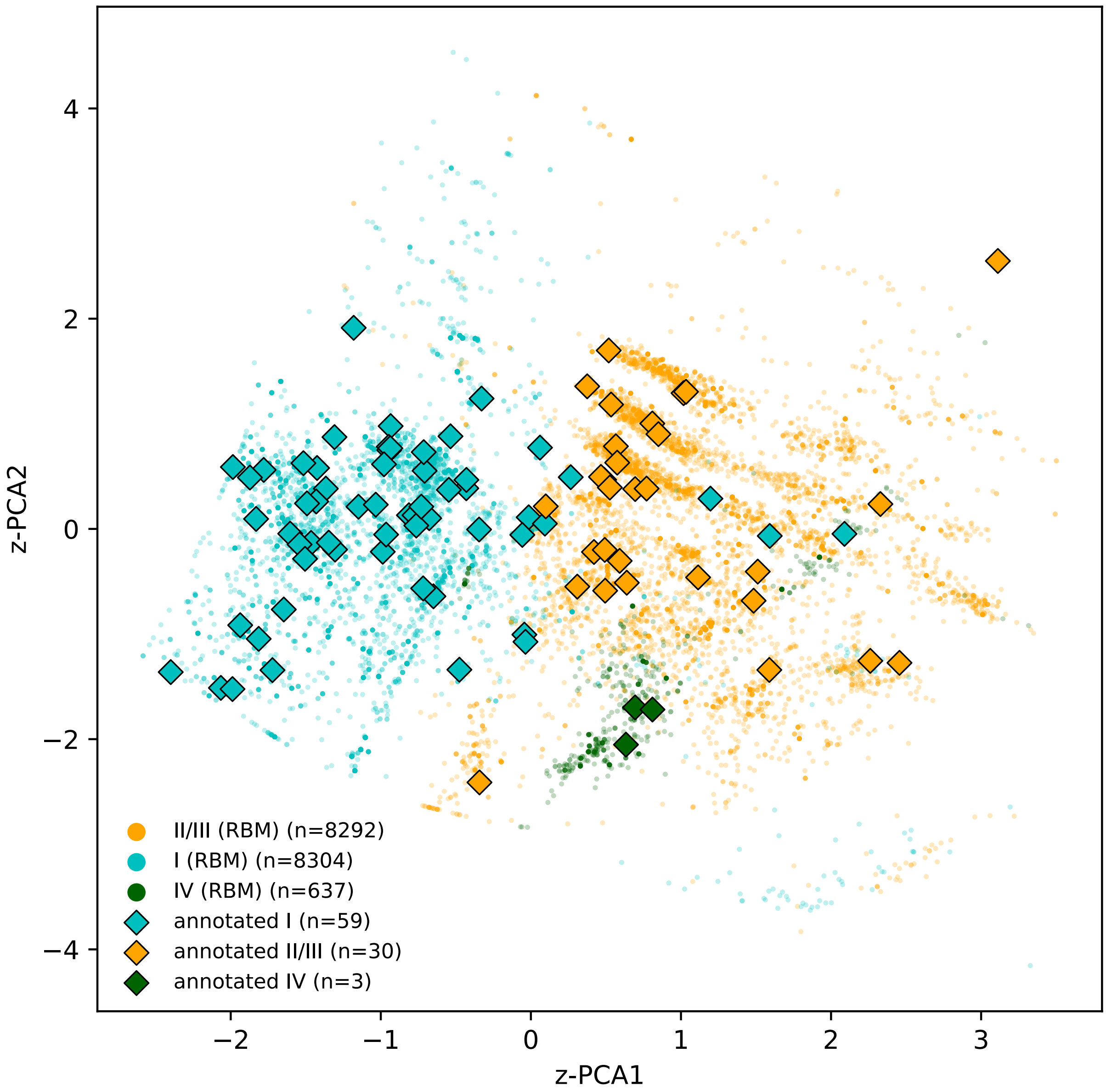


Suppl. Figure 8: Clustering of WW domain type in the latent space of a variational auto-encoder and comparison with the RBM classification. Experimentally assessed sequences are in diamond.

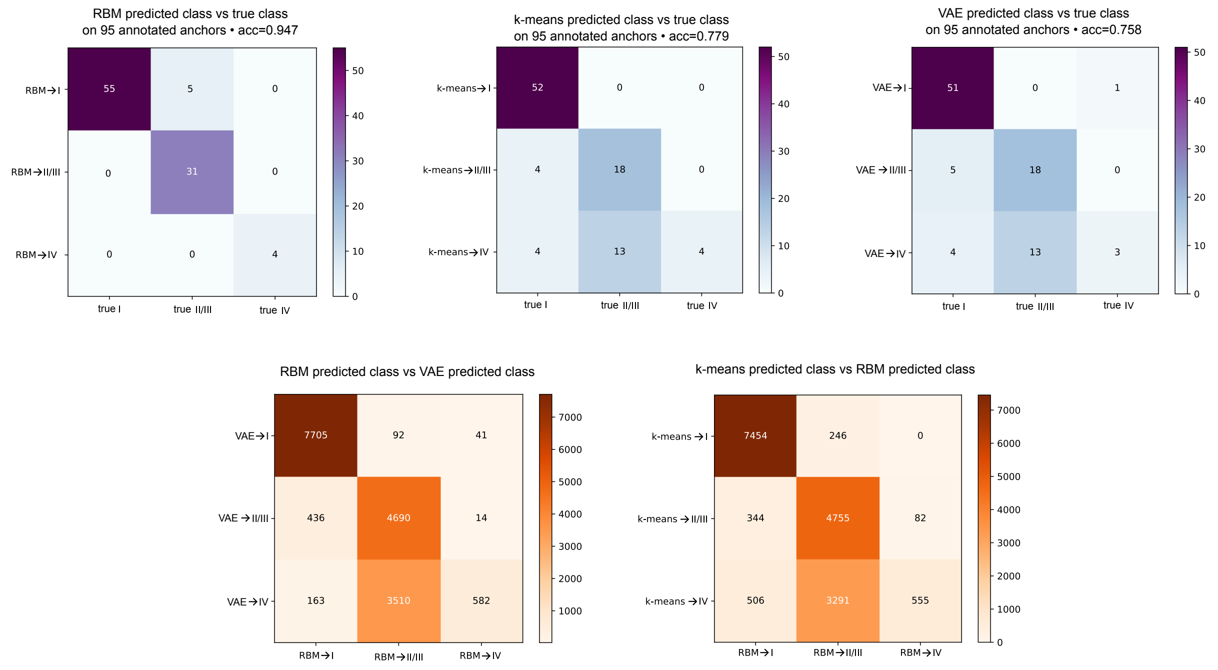


Suppl. Figure 9: Confusion matrices of the classification of the WW domain types. In blue, the three methods of prediction (RBM, sequence $k$-means and variational auto-encoder) are compared with the true type of the domains. In orange, the agreement between the RBM predictions and the other methods are shown.

### B Tested sequences

Tested sequences are given in the tables below, the notation of the sequence gives the name, the class, and minus the log-likelihood of the sequence according to the RBM. All highlighted sequences are natural wild-types.

#### B.1 Tested wild-types

Suppl. Table 1: Tested wild-types sequences and relative responses

| Seq# | notation | Sequence | C1 peptide | C2 peptide | C4 peptide |
|---|---|---|---|---|---|
| seq01 | >wt.01\|classI_s37.73 | LPAGWEMAKTSS-GQRYFLNHIDQTTTWQDP | 0.05±0.014 | 0.03±0.016 | -0.02±0.027 |
| seq02 | >wt.02\|classI_s19.71 | LPPGWERRADSL-GRTYYVDHNTRTTTWTRP | 4.48±0.258 | 0.13±0.026 | 0.04±0.033 |
| seq03 | >wt.03\|classI_s20.15 | LPPGWEQRYTPE-GRPYFVDHNTRTTTWVDP | 3.79±0.551 | 0.22±0.136 | 0.09±0.065 |
| seq04 | >wt.04\|classI_s23.19 | LPPGWERRTDNF-GRTYYVDHNTRTTTWKRP | 4.86±0.351 | 0.37±0.085 | 0.07±0.061 |
| seq05 | >wt.05\|classI_s24.53 | LPSGWEERKDAK-GRTYYVNHNNRTTTWTRP | 14.7±1.849 | 0.5±0.232 | -0.32±0.133 |
| seq06 | >wt.06\|classI_s24.64 | LPSGWEQRFTPE-GRAYFVDHNTRTTTWVDP | 5.56±0.41 | 0.12±0.085 | -0.05±0.07 |
| seq07 | >wt.07\|classI_s24.89 | LPPGWEMKYTSE-GVRYFVDHNTRTTTFKDP | -0.16±0.045 | -0.02±0.027 | -0.15±0.041 |
| seq08 | >wt.08\|classI_s25.83 | LPPGWEIRKDGR-GRVYYVDHNTRKTTWQRP | 4.96±0.187 | 0.39±0.06 | -0.01±0.035 |
| seq09 | >wt.09\|classI_s26.01 | LPPGWEERTHTD-GRVFFINHNIKKTQWEDP | -0.08±0.105 | 0.04±0.128 | -0.07±0.094 |
| seq10 | >wt.10\|classI_s26.29 | LPPGWEKRTDSN-GRVYFVNHNTRITQWEDP | 1.36±0.281 | 0.03±0.062 | -0.1±0.042 |
| seq11 | >wt.11\|classI_s16.87 | LPPGWERRVDSL-GRTYYVDHNTRTTTWTRP | 8.87±0.439 | 0.09±0.041 | 0.01±0.043 |
| seq12 | >wt.12\|classII | VILPPGWQSYLSPQGRRYYVNTTTNETTWERPSSSPG | 0 | 0.1 | - |
| seq13 | >wt.13\|classII | ATAVSEWTEYKTADGKTYYYNNRTLESTWEKPQELKE | 3 | 5 | - |
| seq14 | >wt.14\|classII_s75.12 | GIEMGDWQEVWDENTGCYYYWNTQTNEVTWELPQYLATQ | 0.02±0.005 | 0.92±0.082 | 0±0.007 |
| seq15 | >wt.15\|classII | SGAKSMWTEHKSPDGRTYYYNTETKQSTWEKPDDLKT | 8.7 | 16.7 | - |
| seq16 | >wt.16\|classII | YDSADDWSEHISSSGKKYYYNCRTEVSQWEKPKEWLER | 0.4 | 3.7 | - |
| seq17 | >wt.17\|classII | SDLPAGWMRVQDTSGTYYWHIPTGTTQWEPPGRASP | 0 | 0.1 | - |
| seq18 | >wt.18\|classII | DLPPGWKRVSDIAGTYYWHIPTGTTQWERPVS | 0 | 0 | - |
| seq19 | >wt.19\|classII | AGLPPGWRKIHDAAGTYYWHVPSGSTQWQRPTWELGD | 0 | 0 | - |
| seq20 | >wt.20\|classII | GPPRALWSEHVAPDGRIYYYNADDKQSVWEKPSVLKSK | 0.1 | 1.2 | - |
| seq21 | >wt.21\|classII | NATPKGWSCHWDRDHRRYFYVNEQSGESQWEFPDGEEE | 0 | 3.3 | - |
| seq22 | >wt.22\|classII | APLPGEWKPCQDITGDIYYFNFANGQSMWDHPCDEHY | 0.4 | 0 | - |
| seq23 | >wt.23\|classII | IVLPPNWKTARDPEGKIYYYHVITRQTQWDPPTWESP | 0.2 | 0.5 | - |
| seq24 | >wt.24\|classII | DPSKGRWVEGITSEGYHYYYDLISGASQWEKPEGFQGD | 0.1 | 0.4 | - |
| seq25 | >wt.25\|classII | TAVKTVWVEGLSEDGFTYYYNTETGESRWEKPDDFIPH | 0.2 | 0 | - |
| seq26 | >wt.26\|classII | VSSDGSWLKLNLHKKYDYYYNTDSKESSWVTPESCFYK | 0.2 | 0 | - |
| seq27 | >wt.27\|classIV_s32.36 | LPPGWEKRMSRSSGRVYYFNHITNASQWERP | 0.06±0.01 | 0.12±0.008 | 1.46±0.201 |
| seq42 | >B2.1\|C1C1\|18_s23.98 | LPEGWEIRYTRE-GVRYFVDHNTRTTTFKDP | 26.72±2.737 | 0.03±0.017 | 0±0.015 |

In red are sequences that are repeated in more than one path.

#### B.2 1st batch

First batch of tested sequences. Parameters: $\Lambda = 0.1, \beta = 3, T = 27$.

Suppl. Table 2: First-batch tested sequences and relative responses. The columns give i) the sequence number, ii) the notation indicating: the Batch (B), the specificities XY of the anchoring sequences (CXCY) the index along the designed path starting from 0, and the RBM global score iii) the Sequence, iv) the experimental responses to the peptides C1,C2,C3.

| Seq# | notation | Sequence | C1 peptide | C2 peptide | C4 peptide |
|---|---|---|---|---|---|
| seq01 | >B1\|C1C4\|0_s37.73 | LPAGWEMAKTSS-GQRYFLNHIDQTTTWQDP | 0.05±0.014 | 0.03±0.016 | -0.02±0.027 |
| seq28 | >B1\|C1C4\|5_s28.99 | LPAGWEMRKTSS-GRRYFVNHITRTTTWQDP | 4.48±0.495 | 0.1±0.012 | 0±0.018 |
| seq29 | >B1\|C1C4\|10_s24.11 | LPAGWEMRRTPS-GRVYFVNHITRTTQWEDP | 6.86±0.239 | 0.02±0.029 | -0.02±0.033 |
| seq30 | >B1\|C1C4\|12_s22.26 | LPPGWEERRTPS-GRVYFVNHITRTTQWEDP | 7.77±0.363 | 0±0.062 | -0.06±0.075 |
| seq31 | >B1\|C1C4\|15_s19.74 | LPPGWEERRDPS-GRVYYVNHITRTTQWERP | 8.88±0.959 | 0.01±0.043 | -0.04±0.047 |
| seq32 | >B1\|C1C4\|17_s19.25 | LPPGWEERVDPS-GRVYYVNHITRTTQWERP | 9.48±0.179 | 0.02±0.028 | -0.03±0.048 |
| seq33 | >B1\|C1C4\|19_s20.59 | LPPGWEERVSRS-GRVYYVNHITRTTQWERP | 7.9±0.919 | 0.47±0.053 | 0±0.019 |
| seq34 | >B1\|C1C4\|21_s23.0 | LPPGWEKRMSRS-GRVYYVNHITRTTQWERP | 5.76±0.339 | 0.52±0.024 | -0.01±0.016 |
| seq35 | >B1\|C1C4\|24_s29.01 | LPPGWEKRMSRSSGRVYYVNHITRASQWERP | 1.97±0.117 | 0.1±0.018 | 0.11±0.021 |
| seq27 | >B1\|C1C4\|26_s32.36 | LPPGWEKRMSRSSGRVYYFNHITNASQWERP | 0.06±0.01 | 0.12±0.008 | 1.46±0.201 |

This I-IV path is represented in Suppl. Figure SI10A; as is clear from Table 2 the wild-type of Class I (Yap1) has a low RBM score and was not responsive. Apart from the wild-type the designed path is always functional and responsive to C1 and C4 peptides for the last sequence. Seq.33 and 34 have also a mild response to the C2 peptide showing some sign of promisquity.

#### B.3 2nd batch

Second batch of tested sequences. Parameters: $\Lambda = 0.1, \beta = 3$.

The I-I path is represented in main Figure 2.

The I-IV path is represented in Suppl. Figure SI10B. Seq.34 and 35 are also found in the C1-C4 paths of batch1 and seq.34 has, as already stressed, some response to C2 peptide. Seq.48 shows a response both to C1 peptide and to C4 peptide.

Suppl. Table 3: Second batch tested sequences and relative responses, Path B2.1 (with notation highlighted in green) is shown in Figure 2 of the main text. The columns give i) the sequence number, ii) the notation indicating: the Batch (B), the specificities XY of the anchoring sequences (CXCY) the index along the designed path starting from 0 (corresponding for path B2.1 to the numbers in the main figure 2 minus 1), and the RBM global score iii) the Sequence, iv) the experimental responses to the peptides C1,C2,C3.

| Seq# | notation | Sequence | C1 peptide | C2 peptide | C4 peptide |
|---|---|---|---|---|---|
| seq02 | >B2.1\|C1C1\|0_s19.71 | LPPGWERRADSL-GRTYYVDHNTRTTTWTRP | 4.48±0.258 | 0.13±0.026 | 0.04±0.033 |
| seq36 | >B2.1\|C1C1\|2_s16.14 | LPPGWERRVDPL-GRTYYVDHNTRTTTWTRP | 7.09±0.284 | 0.21±0.034 | 0.11±0.03 |
| seq37 | >B2.1\|C1C1\|5_s15.94 | LPPGWERRVDPN-GRTYYVDHNTRTTTWTRP | 10±1.184 | 0.21±0.034 | 0.16±0.068 |
| seq38 | >B2.1\|C1C1\|7_s18.68 | LPPGWERRVDPN-GRVYYVDHNTRTTTWTDP | 4.29±0.364 | 0.29±0.022 | 0.19±0.038 |
| seq39 | >B2.1\|C1C1\|10_s19.03 | LPPGWERRYTPN-GRVYFVDHNTRTTTWTDP | 7.44±0.447 | 0.18±0.01 | 0.09±0.028 |
| seq40 | >B2.1\|C1C1\|12_s18.46 | LPPGWEIRYTPE-GRVYFVDHNTRTTTWTDP | 7.29±0.464 | 0.21±0.017 | 0.12±0.022 |
| seq41 | >B2.1\|C1C1\|15_s17.89 | LPPGWEIRYTPE-GVRYFVDHNTRTTTFTDP | 5.58±0.2 | 0.17±0.022 | 0.08±0.045 |
| seq42 | >B2.1\|C1C1\|18_s23.98 | LPEGWEIRYTRE-GVRYFVDHNTRTTTFKDP | 26.72±2.737 | 0.03±0.017 | 0±0.015 |
| seq11 | >B2.2\|C1C4\|1_s16.87 | LPPGWERRVDSL-GRTYYVDHNTRTTTWTRP | 8.87±0.439 | 0.09±0.041 | 0.01±0.043 |
| seq43 | >B2.2\|C1C4\|3_s15.77 | LPPGWERRVDPL-GRTYYVDHNTRTTTWQRP | 8.44±0.61 | 0.07±0.05 | 0.01±0.058 |
| seq44 | >B2.2\|C1C4\|6_s15.98 | LPPGWERRVDPN-GRTYYVDHNTRTTTWERP | 6.8±0.734 | -0.04±0.119 | -0.11±0.126 |
| seq45 | >B2.2\|C1C4\|8_s17.68 | LPPGWEERVDPN-GRTYYVDHNTRTTQWERP | 12.65±1.452 | -0.03±0.108 | -0.13±0.106 |
| seq46 | >B2.2\|C1C4\|11_s18.82 | LPPGWEERVDPS-GRTYYVNHITRTTQWERP | 7.37±0.658 | 0.08±0.052 | -0.01±0.075 |
| seq47 | >B2.2\|C1C4\|13_s19.62 | LPPGWEERVDRS-GRVYYVNHITRTTQWERP | 9.27±0.431 | 0.05±0.042 | -0.04±0.035 |
| seq34 | >B2.2\|C1C4\|16_s22.95 | LPPGWEKRMSRS-GRVYYVNHITRTTQWERP | 5.76±0.339 | 0.52±0.024 | -0.01±0.016 |
| seq35 | >B2.2\|C1C4\|19_s28.96 | LPPGWEKRMSRSSGRVYYVNHITRASQWERP | 1.97±0.117 | 0.1±0.018 | 0.11±0.021 |
| seq48 | >B2.2\|C1C4\|20_s31.00 | LPPGWEKRMSRSSGRVYYVNHITNASQWERP | 0.49±0.083 | 0.12±0.03 | 0.73±0.122 |
| seq27 | >B2.2\|C1C4\|21_s32.31 | LPPGWEKRMSRSSGRVYYFNHITNASQWERP | 0.06±0.01 | 0.12±0.008 | 1.46±0.201 |

#### B.4 3rd batch

Third batch of tested sequences. Parameters: $\Lambda = 0.1$, $\beta = 3$.

Suppl. Table 4: Third batch tested sequences and relative responses, Path B3.1 (with notation highlighted in green) is shown in Figure 3 of the main text. Path B3.2 corresponds to a direct path where starting from the initial sequence only mutations in amino acids of final sequences are possible, see Suppl. Figure SI10C. The columns give i) the sequence number, ii) the notation indicating: the Batch (B), the specificities XY of the anchoring sequences (CXCY) the index along the designed path starting from 0 (Corresponding for path B3.1 to the numbers in Figure 3 minus 1) iii) the Sequence, iv) the experimental responses to the peptides C1,C2,C3.

| Seq# | notation | Sequence | C1 peptide | C2 peptide | C4 peptide |
|---|---|---|---|---|---|
| seq02 | >B3.1\|C1C4\|0_s19.72 | LPPGWERRADSL-GRTYYVDHNTRTTTWTRP | 4.48±0.258 | 0.13±0.026 | 0.04±0.033 |
| seq49 | >B3.1\|C1C4\|4_s15.55 | LPPGWERRVDPR-GRTYYVDHNTRTTTWQRP | 6.8±0.201 | 0.05±0.038 | -0.05±0.075 |
| seq50 | >B3.1\|C1C4\|8_s17.24 | LPPGWEERVDPS-GRTYYVDHNTRTTTWERP | 6.32±0.437 | 0.03±0.098 | -0.11±0.076 |
| seq51 | >B3.1\|C1C4\|11_s19.04 | LPPGWEERVDRS-GRTYYVNHNTRTTQWERP | 10.44±0.521 | 0.13±0.019 | 0.02±0.026 |
| seq47 | >B3.1\|C1C4\|13_s19.64 | LPPGWEERVDRS-GRVYYVNHITRTTQWERP | 8.71±0.546 | 0.09±0.014 | -0.01±0.024 |
| seq52 | >B3.1\|C1C4\|15_s21.59 | LPPGWEKRVSRS-GRVYYVNHITRTTQWERP | 6.21±0.554 | 0.39±0.028 | 0.02±0.036 |
| seq53 | >B3.1\|C1C4\|17_s25.15 | LPPGWEKRMSRS-GRVYYVNHITRTSQWERP | 5.22±0.2 | 0.35±0.01 | 0.02±0.021 |
| seq54 | >B3.1\|C1C4\|18_s27.18 | LPPGWEKRMSRS-GRVYYVNHITRASQWERP | 4.7±0.549 | 0.43±0.065 | 0.05±0.023 |
| seq35 | >B3.1\|C1C4\|19_s28.97 | LPPGWEKRMSRSSGRVYYYVNHITRASQWERP | 1.97±0.117 | 0.1±0.018 | 0.11±0.021 |
| seq48 | >B3.1\|C1C4\|20_s31.02 | LPPGWEKRMSRSSGRVYYYVNHITNASQWERP | 0.49±0.083 | 0.12±0.03 | 0.73±0.122 |
| seq27 | >B3.1\|C1C4\|21_s32.32 | LPPGWEKRMSRSSGRVYYYFNHITNASQWERP | 0.06±0.01 | 0.12±0.008 | 1.46±0.201 |
| seq02 | >B3.2\|C1C4\|0_s19.72 | LPPGWERRADSL-GRTYYVDHNTRTTTWTRP | 4.48±0.258 | 0.13±0.026 | 0.04±0.033 |
| seq55 | >B3.2\|C1C4\|3_s19.84 | LPPGWERRMDSS-GRTYYVDHNTRTTTWERP | 5.02±0.485 | 0.3±0.077 | 0.18±0.068 |
| seq56 | >B3.2\|C1C4\|5_s21.94 | LPPGWERRMDRS-GRTYYVDHNTRTTQWERP | 9.24±0.447 | 0.24±0.057 | 0.14±0.029 |
| seq57 | >B3.2\|C1C4\|7_s22.47 | LPPGWERRMDRS-GRVYYVNHNTRTTQWERP | 4.57±0.394 | 0.28±0.084 | 0.2±0.083 |
| seq58 | >B3.2\|C1C4\|9_s22.90 | LPPGWEKRMDRS-GRVYYVNHITRTTQWERP | 3.87±0.362 | 0.15±0.027 | 0.08±0.026 |
| seq53 | >B3.2\|C1C4\|11_s25.15 | LPPGWEKRMSRS-GRVYYVNHITRTSQWERP | 5.22±0.2 | 0.35±0.01 | 0.02±0.021 |
| seq35 | >B3.2\|C1C4\|13_s28.97 | LPPGWEKRMSRSSGRVYYYVNHITRASQWERP | 1.97±0.117 | 0.1±0.018 | 0.11±0.021 |
| seq48 | >B3.2\|C1C4\|14_s31.02 | LPPGWEKRMSRSSGRVYYYVNHITNASQWERP | 0.49±0.083 | 0.12±0.03 | 0.73±0.122 |
| seq27 | >B3.2\|C1C4\|15_s32.32 | LPPGWEKRMSRSSGRVYYYFNHITNASQWERP | 0.06±0.01 | 0.12±0.008 | 1.46±0.201 |

#### B.5 4th batch

Fourth batch of tested sequences. Parameters: $\Lambda = 0.1, \beta = 3$.

Suppl. Table 5: Fourth batch tested sequences and relative responses, all sequences of batch four were tested with [GI] and [QYLATQ] before and after the sequences mentioned in the table, Path B4.1 (with notation highlighted in green) is shown in Figure 4 of the main text. See Suppl. Figure SI10D for the second path.The columns give i) the sequence number, ii) the notation indicating: the Batch (B), the specificities XY of the anchoring sequences (CXCY) the index along the designed path starting from the designed sequence 80 as the initial part of the designed path, up to sequence 80 , shown in Suppl figure13 has mutations on the same residue with different amino acids (apart from the wild type it corresponds to the numbers in Figure 3 plus 74, notice also that the sequence at step 85 (starting from 0) in the table is the same of the one at step 81 as shown in the full path in Suppl figure13 ) iii) the Sequence, iv) the experimental responses to the peptides C1,C2,C3.

| Seq# | notation | Sequence | C1 peptide | C2 peptide | C4 peptide |
|---|---|---|---|---|---|
| seq02x | >B4.1\|C1C2\|0_s19.70 | GILPPGWERRADSL-GRTYYVDHNTRTTTWTR PQYLATQ | 6.01$\pm$0.229 | 0.16$\pm$0.017 | 0.01$\pm$0.022 |
| seq59 | >B4.1\|C1C2\|80_s15.59 | GILPPGWERRVDPN-GRTYYVDHNTRTTTWQRPQYI | 7.87$\pm$0.297 | 0.18$\pm$0.025 | 0.01$\pm$0.044 |
| seq49 | >B4.1\|C1C2\|85_s15.35 | GILPPGWERRVDPR-GRTYYVDHNTRTTTWQRPQYI | 7.55$\pm$0.314 | 0.13$\pm$0.016 | 0.01$\pm$0.026 |
| seq60 | >B4.1\|C1C2\|90_s16.7 | GILPPGWEERVDPN-GRTYYVDHNTRTTTWQRPQYI | 8.79$\pm$0.798 | 0.09$\pm$0.022 | -0.03$\pm$0.035 |
| seq61 | >B4.1\|C1C2\|95_s25.47 | GILPPGWEERVDPNTGRTYYYVNHQTRETTWERPQY | 11.55$\pm$0.883 | 0.07$\pm$0.042 | -0.06$\pm$0.06 |
| seq84 | >B4.1\|C1C2\|96_s28.34 | GILPPGWEERVDPNTGRTYYYVNHQTNETTWERPQY | 0.22$\pm$0.071 | 0.12$\pm$0.054 | -0.03$\pm$0.023 |
| seq85 | >B4.1\|C1C2\|97_s30.61 | GILPPGWEERVDPNTGRTYYYVNTQTNETTWERPQY | 0.06$\pm$0.033 | 0.13$\pm$0.058 | -0.04$\pm$0.020 |
| seq86 | >B4.1\|C1C2\|98_s31.06 | GILPPGWEERVDPNTGRTYYWNTQTNETTWERPQY | 0.04$\pm$0.023 | 0.14$\pm$0.054 | -0.03$\pm$0.014 |
| seq87 | >B4.1\|C1C2\|99_s30.03 | GILPPGWEEVVDPNTGRTYYWNTQTNETTWERPQY | 0.09$\pm$0.035 | 2.74$\pm$1.139 | -0.01$\pm$0.014 |
| seq62 | >B4.1\|C1C2\|100_s30.45 | GILPPGWQEVVDPNTGRTYYWNTQTNETTWERPQY | 0.09$\pm$0.007 | 1.71$\pm$0.049 | 0$\pm$0.033 |
| seq63 | >B4.1\|C1C2\|105_s34.14 | GILPPGWQEVWDENTGRYYYWNTQTNEVTWELPQ | 0.06$\pm$0.028 | 1.61$\pm$0.051 | -0.01$\pm$0.047 |
| seq14 | >B4.1\|C1C2\|110_s75.12 | GIEMGDWQEVWDENTGCYYYWNTQT NEVTWELPQYLATQ | 0.02$\pm$0.005 | 0.92$\pm$0.082 | 0$\pm$0.007 |
| seq02x | >B4.2\|C1C2\|0_s19.70 | GILPPGWERRADSL-GRTYYVDHNTRTTTWTR PQYLATQ | 6.01$\pm$0.229 | 0.16$\pm$0.017 | 0.01$\pm$0.022 |
| seq59 | >B4.2\|C1C2\|80_s15.59 | GILPPGWERRVDPN-GRTYYVDHNTRTTTWQRPQYI | 7.55$\pm$0.314 | 0.13$\pm$0.016 | 0.01$\pm$0.026 |
| seq65 | >B4.2\|C1C2\|85_s22.36 | GI-P-GWEERVDPN-GRTYYVDHNTRTTTWERPQYLA | 0.2$\pm$0.04 | 0.01$\pm$0.117 | -0.06$\pm$0.217 |
| seq66 | >B4.2\|C1C2\|90_s22.92 | GI----WEERVDPASGRTYYVNHNTRTTTWERPQYL | 1.29$\pm$0.156 | 0.18$\pm$0.081 | 0.1$\pm$0.114 |
| seq67 | >B4.2\|C1C2\|95_s27.75 | GI----WEEVVDPASGRTYYWNTETRETTWERPQYL | 0.08$\pm$0.014 | 1.97$\pm$0.121 | 0.01$\pm$0.025 |
| seq68 | >B4.2\|C1C2\|100_s27.65 | GI----WQEVVDPASGRYYYWNTETNEVTWELPQYL | 0.03$\pm$0.015 | 0.41$\pm$0.009 | 0.01$\pm$0.02 |
| seq69 | >B4.2\|C1C2\|105_s31.75 | GI---DWQEVWDENTGRYYYWNTETNEVTWELPQY | 0.02$\pm$0.005 | 2.39$\pm$0.125 | -0.01$\pm$0.011 |
| seq14 | >B4.2\|C1C2\|110_s75.12 | GIEMGDWQEVWDENTGCYYYWNTQT NEVTWELPQYLATQ | 0.02$\pm$0.005 | 0.92$\pm$0.082 | 0$\pm$0.007 |

#### B.6 Scrambled Paths

Suppl. Table 6: Scrambled paths sequences and relative responses, all sequences of batch four SCR.1 bath were tested with [GI] and [QYLATQ] before and after the sequences mentioned in the table

| Seq# | notation | Sequence | C1 peptide | C2 peptide | C4 peptide |
|---|---|---|---|---|---|
| seq02 | >SCR.1\|C1C2\|1_s19.70 | LPPGWERRADSL-GRTYYVDHNTRTTTWTRP | 6.01±0.229 | 0.16±0.017 | 0.01±0.022 |
| seq70 | >SCR.1\|C1C2\|2 | EMGDWERRADSL-GCTYYVDHNTRTTTWTRP | 0±0.001 | 0±0.001 | 0±0.001 |
| seq71 | >SCR.1\|C1C2\|3 | EMGDWERRVDPL-GCYYYVDHNTRTVTWTLP | 0±0.001 | 0±0.001 | 0±0.001 |
| seq72 | >SCR.1\|C1C2\|4 | EMGDWQRVVDPL-GCYYYWDTNTNTVTWTLP | 0.01±0.01 | 0.01±0.007 | 0.01±0.008 |
| seq73 | >SCR.1\|C1C2\|5 | EMGDWQRVVDPNTGCYYYWNTQTNEVTWQLP | 0.01±0.01 | 0.02±0.009 | -0.01±0.007 |
| seq74 | >SCR.1\|C1C2\|6 | EMGDWQEVVDPRTGCYYYWNTQTNEVTWQLP | 0±0.005 | 0.01±0.005 | -0.01±0.005 |
| seq75 | >SCR.1\|C1C2\|7 | EMGDWQEVVDPNTGCYYYWNTQTNEVTWQLP | 0±0.005 | 0.23±0.057 | 0±0.006 |
| seq14 | >SCR.1\|C1C4\|8_s75.12 | EMGDWQEVWDENTGCYYYWNTQTNEVTWELP | 0.02±0.005 | 0.92±0.082 | 0±0.007 |
| seq02 | >SCR.2\|C1C4\|1_s19.72 | LPPGWERRADSL-GRTYYVDHNTRTTTWTRP | 4.48±0.258 | 0.13±0.026 | 0.04±0.033 |
| seq76 | >SCR.2\|C1C4\|2 | LPPGWERRADSL-GRTYYFDHNTNTTTWTRP | 0±0.032 | 0±0.028 | -0.05±0.039 |
| seq77 | >SCR.2\|C1C4\|3 | LPPGWERRADSLSGRTYYFDHNTNATTWTRP | 0.04±0.027 | 0.04±0.034 | -0.02±0.035 |
| seq78 | >SCR.2\|C1C4\|4 | LPPGWERRASSLSGRTYYFDHNTNASTWTRP | 0.05±0.015 | 0.07±0.023 | 0.01±0.028 |
| seq79 | >SCR.2\|C1C4\|5 | LPPGWEKRASSLSGRTYYFDHITNASTWTRP | 0.01±0.012 | 0.03±0.012 | -0.02±0.018 |
| seq80 | >SCR.2\|C1C4\|6 | LPPGWEKRASSLSGRVYYFNHITNASTWTRP | 0.01±0.016 | 0.02±0.011 | -0.02±0.019 |
| seq81 | >SCR.2\|C1C4\|7 | LPPGWEKRASRLSGRVYYFNHITNASQWTRP | 0.03±0.004 | 0.02±0.015 | -0.01±0.013 |
| seq27 | >SCR.2\|C1C4\|8_s32.32 | LPPGWEKRMSRSSGRVYYFNHITNASQWERP | 0.06±0.01 | 0.12±0.008 | 1.46±0.201 |

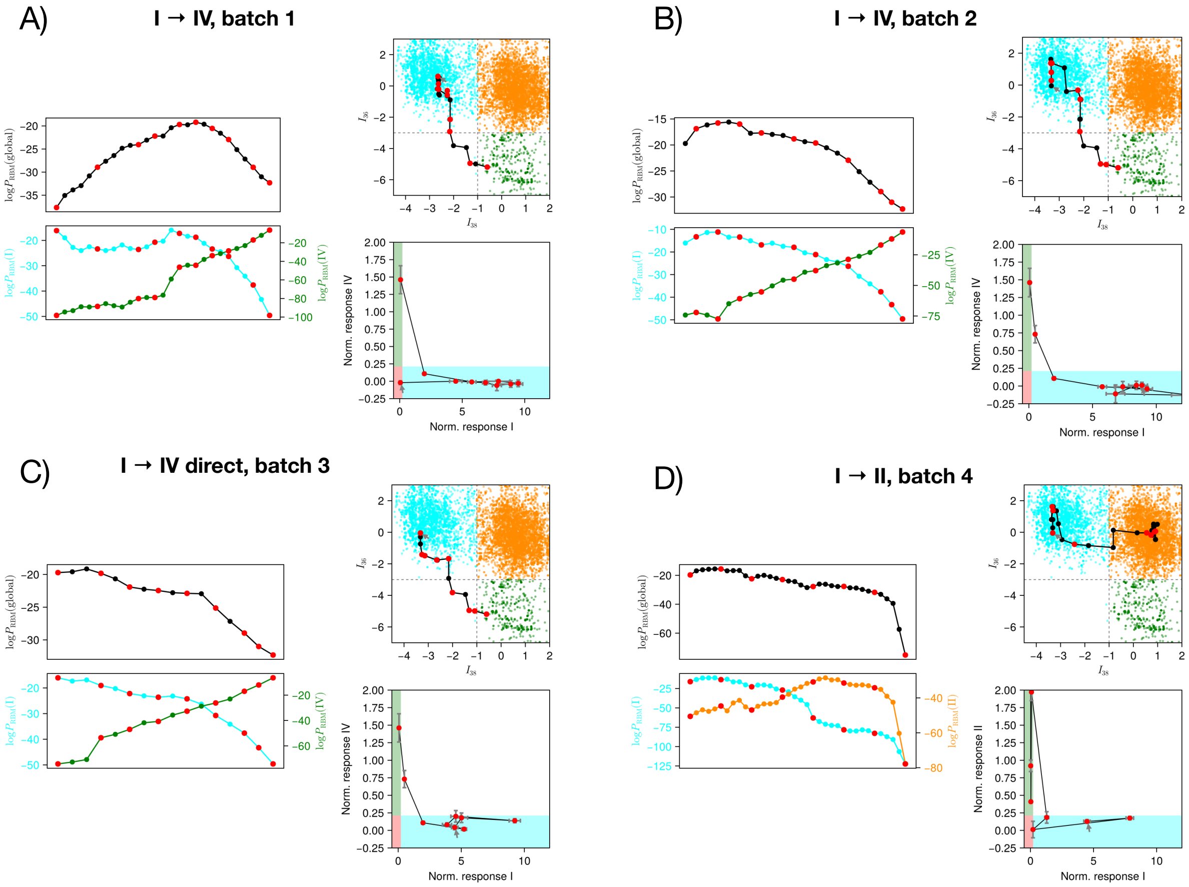


Suppl. Figure 10: Supplementary paths. A) Path I→IV, batch 1. B) Path I→IV, batch 2. C) Direct path I→IV. D) Path I→II, batch 4.

#### B.7 Full generated paths

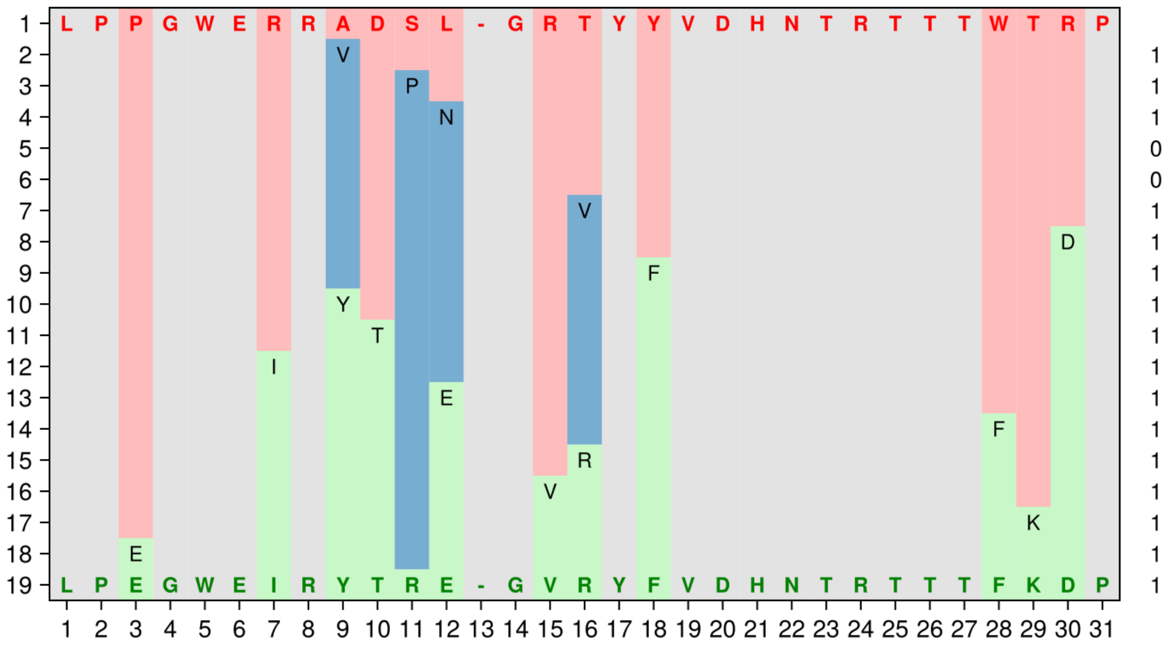


Suppl. Figure 11: Full generated path $I \rightarrow I$.

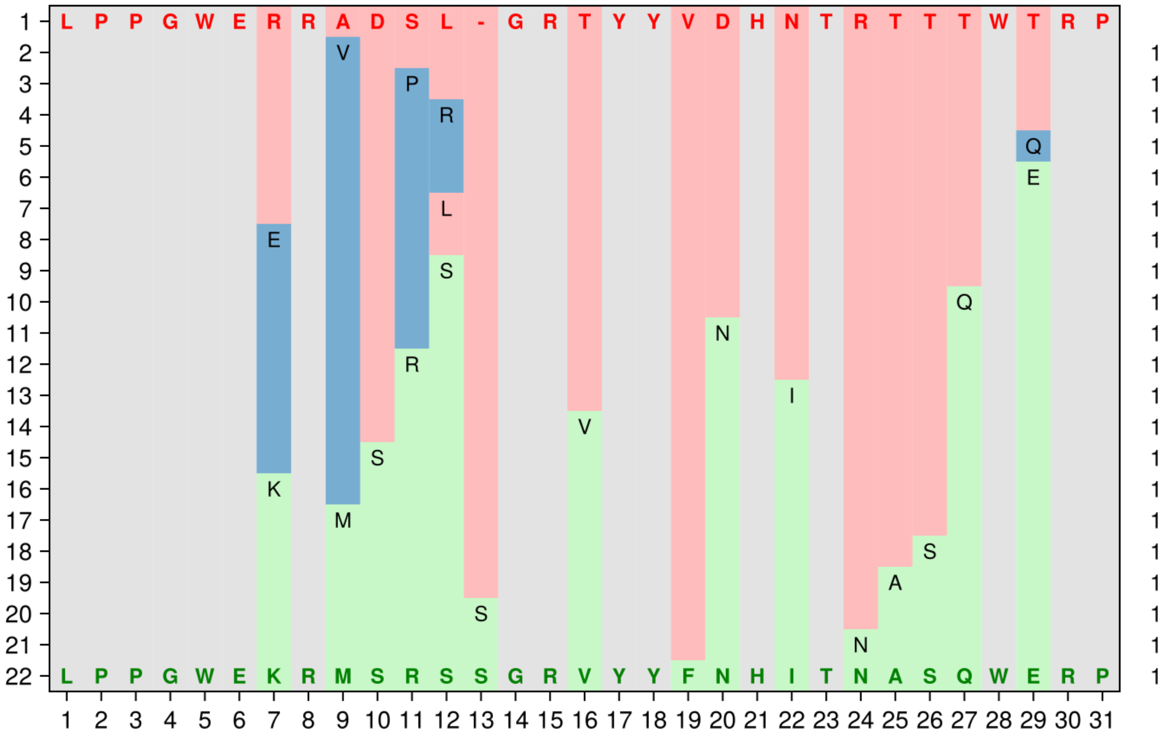


Suppl. Figure 12: Full generated path $I \rightarrow IV$.

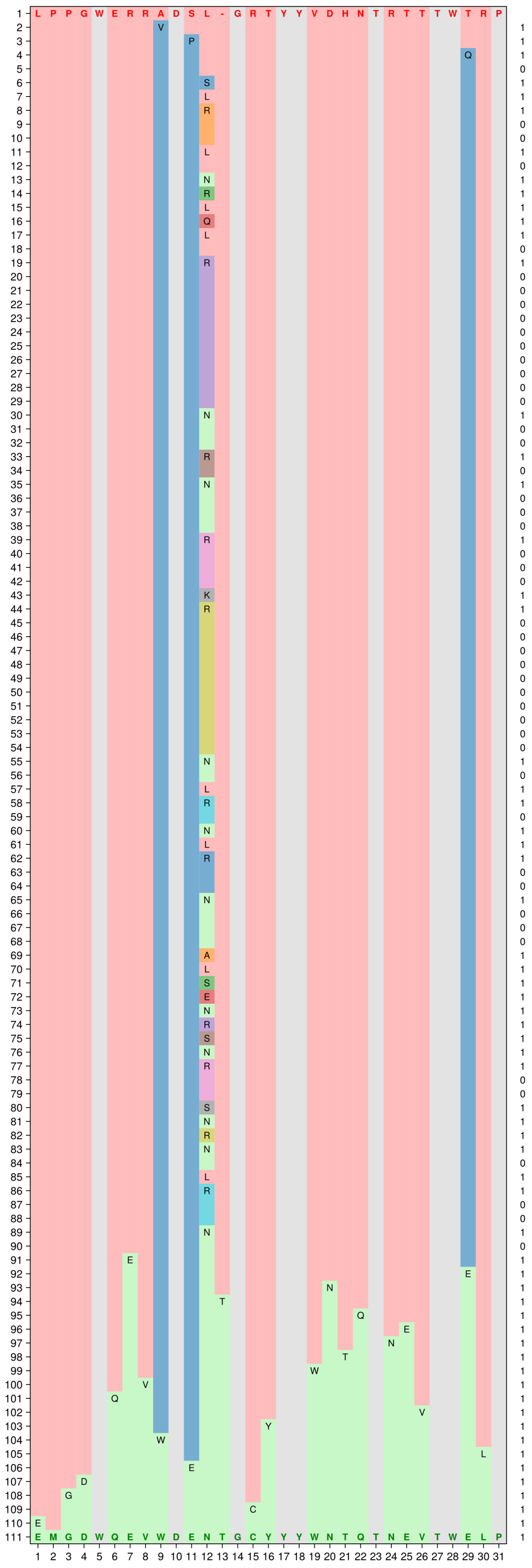

Suppl. Figure 13: Full generated path $I \rightarrow II$.

#### B.8 Ancestral protein reconstruction paths

Suppl. Table 7: Full ASR path $I \to II$. Scores are given according to the global and class-specific RBMs. Time is the time of divergence (average number of mutation per site in the LG+G4 model) separating a sequence with its direct ancestors.

| Seq# | Sequence | $E_{glob}$ | $E_{classI}$ | $E_{classII}$ | $E_{classIV}$ | time |
|---|---|---|---|---|---|---|
| asrseq1 | LPPGWERRADSL-GRTYYVDHNTRTTTWTRP | -19.71 | -16.06 | -50.58 | -74.23 | 0.0 |
| asrseq2 | LPPGWERRVDNL-GRTYYVDHNTRTTTWQRP | -17.61 | -10.63 | -50.48 | -76.55 | 0.07 |
| asrseq3 | LPPGWEARVDQY-GRTYYVDHNTRTTTWQRP | -22.01 | -18.59 | -49.49 | -72.24 | 0.07 |
| asrseq4 | LPPGWEARVDQY-GRTYYVDHNTRTTTWERP | -22.01 | -18.89 | -47.90 | -68.22 | 0.02 |
| asrseq5 | LPPGWEERVDPY-GRTYYVDHNTRTTTWERP | -19.68 | -17.42 | -45.38 | -70.69 | 0.06 |
| asrseq6 | LPPGWEERVDPS-GRTYYVDHNTRTTTWERP | -17.23 | -16.82 | -40.98 | -53.68 | 0.02 |
| asrseq7 | LPPGWEERVDPS-GRTYYVNHNTRTTTWERP | -18.85 | -18.94 | -38.48 | -48.36 | 0.04 |
| asrseq8 | LPPGWEERVDPS-GRTYYVNHTTRTTQWERP | -19.07 | -18.82 | -36.01 | -44.64 | 0.04 |
| asrseq9 | LPPGWEERVDPSSGRTYYVNHTTRTTQWERP | -20.76 | -21.44 | -36.71 | -40.57 | 0.02 |
| asrseq10 | LPPGWKEAVDPSSGRTYYYNTTTRTTQWERP | -31.46 | -58.005 | -33.41 | -45.425 | 0.14 |
| asrseq11 | LPPNWKEAVDPSSGRTYYYNTKTRTTQWERP | -32.29 | -63.59 | -33.74 | -46.02 | 0.05 |
| asrseq12 | LPPNWKEATDPNSGRTYYYNTKTRETTWEKP | -33.67 | -70.17 | -33.86 | -51.65 | 0.10 |
| asrseq13 | LPPNWKEYTTPD-GRKYYYNTQTKETTWEKP | -28.725 | -70.24 | -28.69 | -84.91 | 0.20 |
| asrseq14 | LPPNWKEYTTPD-GRKYYYNTQTKESTWEKP | -29.59 | -73.89 | -28.81 | -80.35 | 0.04 |
| asrseq15 | SQSNWKEYTTDD-GRKYYYNTQTKESTWEKP | -51.34 | -99.96 | -50.97 | -100.96 | 0.11 |
| asrseq16 | SQSQWKEYTTDD-GRPYYYNTQTKESRWEKP | -54.16 | -105.43 | -54.55 | -104.24 | 0.08 |
| asrseq17 | SQSQWKEYTSDD-GRPYYYNTQTKESRWEKP | -54.99 | -106.77 | -54.33 | -87.10 | 0.03 |
| asrseq18 | SQGQWKEYTSDD-GRPYYYNTLTKESRWEKP | -57.26 | -107.58 | -56.33 | -88.03 | 0.04 |
| asrseq19 | SQGQWKEYTSDD-GRPYYYNTLTKETRWEKP | -57.49 | -104.30 | -57.56 | -93.05 | 0.04 |
| asrseq20 | SQGQWKEYMSDD-GRPYYYNTLTKETQWEKP | -57.86 | -98.95 | -59.24 | -90.09 | 0.05 |
| asrseq21 | SQGPWKEYFSDD-GRPYYYNTLTGETQWEKP | -55.13 | -99.77 | -56.89 | -87.63 | 0.11 |
| asrseq22 | AAGPWKEYFSDD-GRPYYYNTLTGETQWEKP | -54.33 | -99.81 | -56.06 | -87.99 | 0.049 |
| asrseq23 | AAGPWKEYWDDE-GRPYYYNTVTGETQWEKP | -56.49 | -102.25 | -57.20 | -77.86 | 0.10 |
| asrseq24 | AAGPWKEYWDDE-GRTYYYNTVTGETQWEKP | -57.11 | -102.04 | -57.09 | -75.83 | 0.03 |
| asrseq25 | AAGDWKEYWDDE-GRTYYYNTVTGETQWEKP | -57.41 | -102.42 | -58.16 | -74.69 | 0.03 |
| asrseq26 | SAGDWQEYWDDE-GRIYYYNTQTGETSWEPP | -60.75 | -101.87 | -60.27 | -80.35 | 0.22 |
| asrseq27 | SMGDWQEYWDEESGRFYYYNTQTGETSWEPP | -73.85 | -117.14 | -72.48 | -90.84 | 0.07 |
| asrseq28 | EMGDWQEVWDENTGCYYYWNTQTNEVTWELP | -75.13 | -122.40 | -77.85 | -101.64 | 0.44 |

Suppl. Table 8: Full ASR path $I \to IV$. Scores are given according to the global and class-specific RBMs. Time is the time of divergence (average number of mutation per site in the LG+G4 model) separating a sequence with its direct ancestors.

| Seq# | Sequence | $E_{glob}$ | $E_{classI}$ | $E_{classII}$ | $E_{classIV}$ | time |
|---|---|---|---|---|---|---|
| asrseq1 | LPPGWERRADSL-GRTYYVDHNTRTTTWTRP | -19.71 | -16.06 | -50.58 | -74.23 | 0.0 |
| asrseq2 | LPPGWERRVDNL-GRTYYVDHNTRTTTWQRP | -17.61 | -10.63 | -50.48 | -76.55 | 0.07 |
| asrseq3 | LPPGWEARVDQY-GRTYYVDHNTRTTTWQRP | -22.01 | -18.59 | -49.49 | -72.24 | 0.07 |
| asrseq4 | LPPGWEARVDQY-GRTYYVDHNTRTTTWERP | -22.01 | -18.89 | -47.90 | -68.22 | 0.02 |
| asrseq5 | LPPGWEERVDPY-GRTYYVDHNTRTTTWERP | -19.68 | -17.42 | -45.38 | -70.69 | 0.06 |
| asrseq6 | LPPGWEERVDPS-GRTYYVDHNTRTTTWERP | -17.23 | -16.82 | -40.98 | -53.68 | 0.02 |
| asrseq7 | LPPGWEERVDPS-GRTYYVNHNTRTTTWERP | -18.85 | -18.94 | -38.48 | -48.36 | 0.04 |
| asrseq8 | LPPGWEERVDPS-GRTYYVNHTTRTTQWERP | -19.07 | -18.82 | -36.01 | -44.64 | 0.04 |
| asrseq9 | LPPGWEERVTPS-GRTYYVNHTTRTTQWERP | -21.26 | -19.64 | -37.25 | -57.37 | 0.02 |
| asrseq10 | LPPGWEERVTPS-GRTYYFNHTTRTSQWERP | -29.23 | -30.21 | -37.55 | -54.71 | 0.07 |
| asrseq11 | LPPGWEERVSRSTGRTYYLNHTTKASQWERP | -27.95 | -39.88 | -43.99 | -26.93 | 0.096 |
| asrseq12 | LPPGWEKRVSRSTGRTYYLNHTTKASQWERP | -28.28 | -41.81 | -45.11 | -23.35 | 0.03 |
| asrseq13 | LPPGWEKRMSRSTGRVYYFNHTTNASQWERP | -31.45 | -50.02 | -44.72 | -15.34 | 0.10 |
| asrseq14 | LPPGWEKRMSRSSGRVYYFNHITNASQWERP | -32.31 | -49.61 | -44.90 | -6.37 | 0.04 |

### C Experimental Protocol

#### C.1 Linker and Beads preparation

In order to produce spectrally distinguishable beads with DBCO functional groups, we used fluorescently modified oligonucleotide linkers. As illustrated in Suppl. Figure 14A, a forward oligonucleotide was designed and ordered with a 5' biotin modification, 4A bases, 25 bases with GC=64%, and a Dibenzocyclooctyne (DBCO) modification at the 3' end. Additionally, three reverse oligonucleotides have been ordered with a 5' modification of either Cy3B, Pacific Blue, or 6-Fam, and one reverse oligonucleotide has been ordered without any modification. The hybridization of the forward oligonucleotide with different ratios of the four reverse oligonucleotides can allow for the generation of eight distinct colors. In our experiments, we utilize seven colors for the DBCO Linkers and reserve no-color beads for Protein A beads coated with anti-SNAP antibody, which we employ for data normalization based on the expression assessment.

To minimized photobleaching of the fluorescent dyes, all handling of the DNA Linker or labeled proteins were conducted in low-light conditions and within light-protection tubes.

Following the mixing of the oligonucleotides in accordance with the ratios explained in Suppl. Figure 14A, an optional denaturation and hybridization step may be conducted to ensure the complete hybridization of the oligonucleotides. However, it should be noted that the oligonucleotides can hybridize at room temperature without denaturation, given that the GC content is high and no secondary structure can be formed at room temperature. All oligonucleotides have been purified by HPLC and resuspended in nuclease-free water to achieve a final concentration of 100 µM for the single-stranded DNA oligos. Following the mixing of the forward and reverse oligonucleotides, a 50 µM solution of the double-stranded DNA linker is obtained.

For each color of beads, 10 µL of Streptavidin magnetic beads (10 mg/ml) were transferred to a new tube, resuspended in 200 µL of wash buffer, and vortexed to ensure uniform suspension. Subsequently, the beads were placed on a magnetic stand for one minute to remove the preservative solution. The supernatant was discarded, and the washing procedure was repeated twice. The DNA Linker for each color was prepared by resuspending it in 1X phosphate-buffered saline buffer (DPBS) with 0.5% Tween-20 (wash buffer), and its concentration was adjusted to 2 µM. Subsequently, 25 µL of the DNA Linker was added to the prewashed streptavidin magnetic beads, and the mixture was agitated gently to prevent precipitation. The beads were incubated with the DNA at 37°C for 30 minutes with continuous mixing at 800 rpm in a thermomixer. Following the incubation period, the tube was placed on the magnetic stand, and the supernatant was removed. The beads were then washed three times with the wash buffer by resuspending them and vortexing gently before returning them to the magnetic stand for separation and buffer removal. Once coated with the DNA Linker, the beads were ready to be resuspended in the peptide solution.

Peptides were synthesized with an N-terminal azide (-N3) group and purified by high-performance liquid chromatography (HPLC), resulting in >95% pureness. Following a solubility test, the peptides were resuspended in an optimal solvent, as advised by the manufacturer, to a final concentration of 4 mg/ml and aliquoted for storage at -80 °C. The peptide solutions were prepared by diluting the peptides in wash buffer to a concentration of 200 µM. The beads coated with the DNA linker were resuspended in 25 µL of peptide solution and vortexed gently to ensure uniform suspension. The beads were incubated with the peptide solution for 16 hours at room temperature with continuous mixing in a tube rotator. Following the incubation period, the tube was placed on the magnetic stand, and the supernatant was removed. The beads were then washed three times with wash buffer, with each wash involving the resuspension of the beads and gentle vortexing before returning them to the magnetic stand for separation and buffer removal. The beads were subsequently resuspended in 100 µL of binding buffer (Tris-HCl 50 mM pH 7.5, EDTA 10 mM,DTT 1 mM and 0.5 % Tween 20). Once coated with the azide peptide, the beads were ready for mixing with different colors.

For the normalization of expression beads, 3.3 µL of Protein A magnetic beads (30 mg/ml) were transferred to a new tube, resuspended in 200 µL of wash buffer, and vortexed to ensure uniform suspension. Subsequently, the beads were placed on a magnetic stand for one minute to remove the preservative solution. The supernatant was then discarded, and the washing procedure was repeated two times. The polyclonal anti-SNAP antibody was prepared by diluting it in wash buffer, and its concentration was adjusted to 1 µM. Subsequently, 20 µL of the diluted antibody was added to the prewashed Protein A magnetic beads, and the mixture was vortexed gently to facilitate binding. The beads were incubated with the antibody at 37°C for 30 minutes with continuous mixing at 800 rpm in a thermomixer. Following the incubation period, the tube was placed on the magnetic stand, and the supernatant was removed.

The beads were then washed three times with wash buffer by resuspending them and vortexing before returning them to the magnetic stand for separation and buffer removal. The beads were resuspended in 100 µl binding buffer. After coating with the anti-SNAP antibody, the beads were ready for mixing with different peptide-coated beads.

The eight prepared beads were combined in equal volumes to create the Beads Mixture that was used during the binding step. Prior to this, an experiment was conducted to ascertain the feasibility of demultiplexing the eight beads in the flow cytometer. We could differentiate between the eight colors.

To validate the design of the multicolored beads system, streptavidin magnetic beads were coated with different peptides using the 7 colors linkers system, and no-color Protein A beads were coated with anti-SNAP antibody. After mixing of all the beads, 128 samples were acquired in the flow cytometer and demultiplexed as explained in Suppl. Figure 14B. In the plot A, the separation of beads is based on their size (Forward Scatter, y-axis) and the absence (gated to plot B) or presence (gated to plot B) of the 6-FAM dye (Green-B Fluorescence, x-axis). In plots B and C, the beads are separated based on the absence or presence of the Cy3B dye (Yellow-B Fluorescence, y-axis in plot A and x-axis in plot C) and the Pacific Blue dye (Yellow-B Fluorescence, x-axis in plot B and y-axis in plot C). This process enables the demultiplexing of eight distinct populations of color-coded beads. An analysis of the percentage for each demultiplexed color was conducted and is shown in Suppl. Figure 14C. The results demonstrate an efficient demultiplexing of the eight colors with minimal variation in the percentage of each color.

To validate the expression normalization system, two experiments were performed using SNAP-tag® Purified Protein From (New England Biolabs, P9312S) as shown in Suppl. Figure 14D. A serial dilution of the SNAP-tag protein is prepared in PBS with SNAP-Surface® Alexa Fluor® 647 dye. The first experiment assesses the binding of the SNAP-tag protein to Protein A beads coated with anti-SNAP antibody, while the second experiment evaluates the binding of the SNAP-tag protein to Protein A beads coated with anti-SNAP antibody as one of eight color-coded beads mixture. All incubation steps were performed at 37°C for 30 minutes. The results in Suppl. Figure 14E showed a strong correlation between the relative concentration and the normalized response indicating that the assay is reproducible and sensitive.

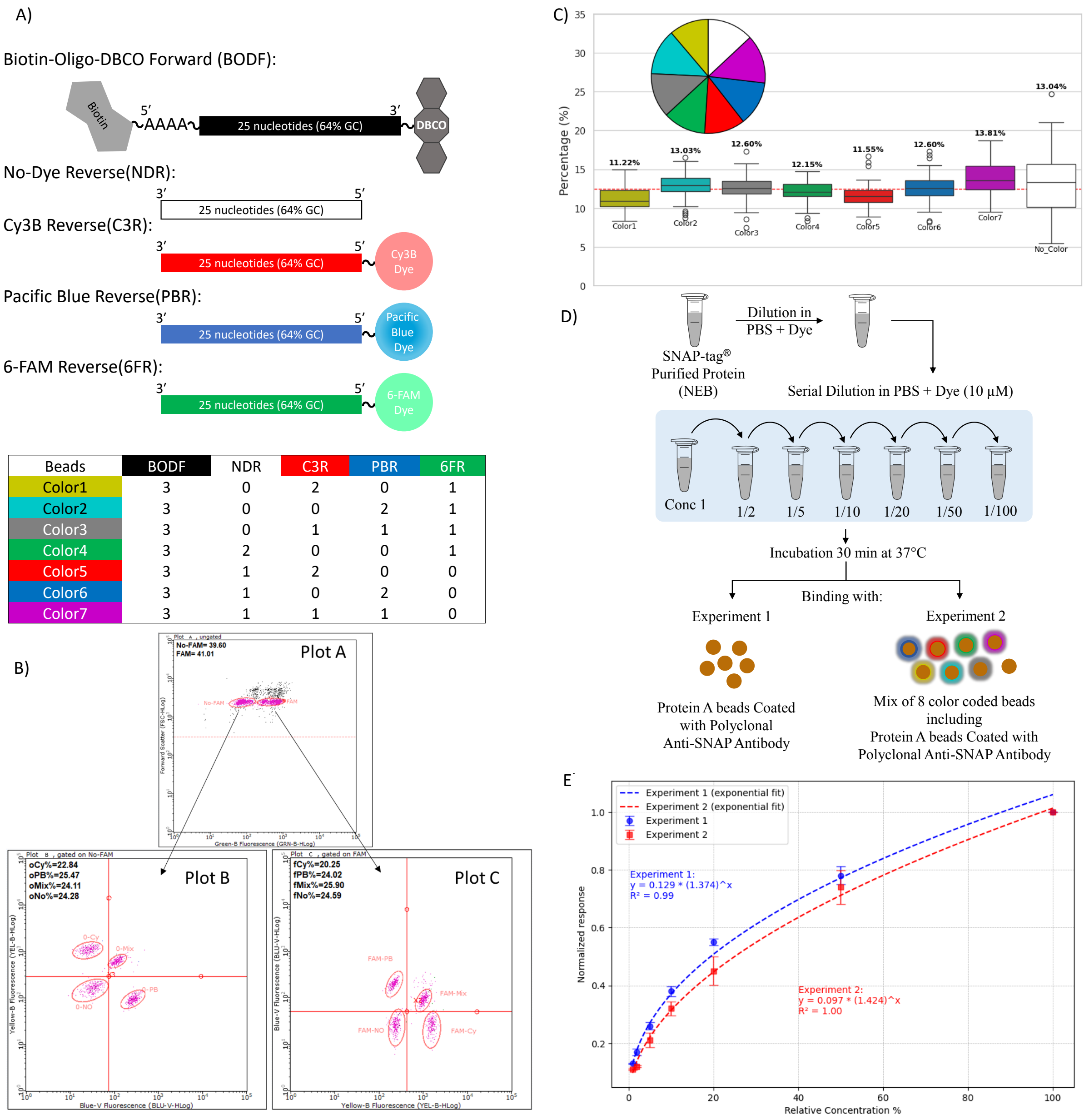


Suppl. Figure 14: Overview of the design, workflow, and validation of bead multiplexing and expression assessment approaches. (A) Schematic representation of oligonucleotide constructs and their fluorescent labeling. A forward oligonucleotide (BODF) of 25 nucleotides with 64% GC content is modified with 5' Biotin group and 3' Dibenzocyclooctyne (DBCO) group. The modified oligonucleotide can be paired with a mixture of four distinct reverse oligonucleotides. The reverse oligonucleotides are designated as: no-dye reverse (NDR), Cy3B reverse (C3R), Pacific Blue reverse (PBR), and 6-FAM reverse (6FR). Each reverse oligonucleotide has a fixed length of 25 nucleotides with 64% GC content. The table illustrates that seven distinct ratios can be employed to generate seven distinguishable colors of DNA linkers for beads labeling (color coding). (B) Flow cytometry plots illustrate the demultiplexing of the color-coded beads. Each bead population shows distinct fluorescence patterns based on their labeling with mixture of reverse oligonucleotides. The gating regions illustrate the effective separation of beads across fluorescence channels. (C) Quantification of bead populations based on fluorescence readout. The pie chart represents the average proportion of each bead population, while the boxplot provides the distribution percentage of fluorescence signals for the individual bead colors in 128 samples acquired in the flow cytometer. (D) Experimental workflow for the validation of expression normalization system (E) The normalized fluorescence intensity data were plotted against the relative concentration percentage. The exponential fitting equations and $R^2$ values for both experiments demonstrate strong agreement between the data sets, indicating that the assay is reproducible and sensitive.

#### C.2 DNA Preparation for Expression

During the validation of the multicolored beads system Suppl. Figure 15A. The DNA sequences encoding the gene to be expressed were designed to be cloned using Gibson Assembly into a plasmid as illustrated in Suppl. Figure 15B. This plasmid contained a T7 RNA Polymerase promoter, a ribosome binding site (RBS), a cloning site, a linkers, SNAP-tag sequence, and a T7 terminator sequence as described in the main text. Following the receipt of eBlocks™ gene fragments at 200 ng for each DNA fragment, all genes were resuspended in 20 µl nuclease-free water, resulting in a final concentration of 10 ng/µl. Twenty nanograms of each gene fragment were combined with 40 ng of the plasmid backbone and 10 µl of 2X Gibson Assembly Master Mix in a 20 µl Gibson Assembly reaction. The reactions were incubated at 50°C for 30 minutes. To validate the assembly step, a no-insert control was performed.

Polymerase chain reaction (PCR) reactions were prepared using 27 µl nuclease-free water, 5 µl ligated plasmids (Gibson assembly reaction), 10 µl of 5X Q5® Reaction Buffer, 2.5 µl of forward primer (10 µM), 2.5 µL of reverse primer (10 µM), 2.5 µL of deoxynucleotide triphosphate (dNTPs at 10 µM), and 0.5 µL of Q5® Hot Start High-Fidelity DNA Polymerase. An initial denaturation step at 98°C for 30 seconds was performed, followed by 25 cycles of denaturation (at 98°C for 15 seconds), annealing (at 68 °C for 15 seconds), and extension (at 72°C for 60 seconds). Subsequently, a final extension step was conducted (at 72 °C for 2 minutes) following the 25 PCR cycles.

Purification of the PCR products was conducted using SPRIselect magnetic beads. This was achieved by mixing equal volumes of the PCR product and the beads, the mixtures were vortexed gently to facilitate DNA binding to the beads, the beads were incubated with the DNA at room temperature for five minutes. Following the incubation period, the tubes were placed on the magnetic stand for one minute, after which the supernatants were removed. The beads were then washed twice with 200 µl of freshly prepared 80% ethanol without resuspension of the beads. The tubes were left open to dry for one minute following the removal of residual ethanol. Subsequently, 10 µl of nuclease-free water were added to the beads, which were then vortexed for one minute. Then, the tubes were placed on the magnetic stand for one minute. The supernatants, which contained the eluted DNA, were then transferred to new tubes. The DNA concentrations were determined using a NanoDrop™ One spectrophotometer from Thermo Scientific. Additionally, agarose gel electrophoresis was conducted to validate the assembly and PCR steps. To ensure the PCR reactions were free of contamination, a no-template control was performed.

#### C.3 Protein Expression and labeling

The purified PCR products were utilized to express the fusion proteins in vitro. Specifically, four microliters of PURExpress Solution (A) were combined with three microliters of PURExpress Solution (B), 0.25 µl of SNAP-Surface® Alexa Fluor® 647 (at 250 µM), 2.75 µl of nuclease-free water, and one microliter of purified PCR products (100 ng/µl). The mixtures were incubated for five minutes at 37°C to initiate transcription, followed by four hours of incubation at 30°C. During this incubation time, protein expression and folding occurred, followed by the formation of a covalent interaction between the expressed SNAP-fusion protein and the fluorescent dye. To achieve optimal fluorescence labeling of the proteins with the SNAP-Surface® Alexa Fluor® 647 dye, an overnight incubation at 4°C was performed. Following the overnight incubation, the labeled proteins were ready for the binding step. A no expression control was performed with no DNA added to the expression mixture to serve as a control during the binding and acquisition steps.

#### C.4 Binding and acquisition in flow cytometer

Binding reactions were performed in 0.2 tubes. Four microliters of bead mixture were added to six microliters of binding buffer and four microliters of labeled protein and mixed by pipetting. Tubes were incubated at 25°C for 30 minutes with continuous mixing at 800 rpm in a thermomixer. After the incubation period, the tubes were placed on the magnetic stand and the supernatants were removed. The beads were then washed twice by resuspension in 200 µL of wash buffer (no vortexing) before being returned to the magnetic stand for separation and buffer removal. The beads were resuspended in 200 µl of wash buffer (filtered through a 0.2 µm filter) and transferred to the flow cytometer plate, as described in Suppl. Figure 15C.

Samples were analyzed using the Guava® easyCyte™ HT system (Merck Millipore). Acquisition settings were configured with the following gain values: Forward Scatter (FSC: 11.8), Side Scatter (SSC: 1.0), Green-B Fluorescence (GRN-B: 5), Blue-V Fluorescence (BLU-V: 5), Yellow-B Fluorescence (YEL-B: 5), Red-R Fluorescence (RED-R: 5), and Near IR-R Fluorescence (NIR-R: 5). Initial gating was performed based on bead size and green fluorescence to exclude larger beads and to distinguish beads containing the 6-FAM dye from those without. Subsequent demultiplexing used blue and yellow fluorescence to identify beads carrying Cy3B dye, Pacific Blue dye, both dyes, or neither dye. Alexa Fluor 647 fluorescence, measured using the Red-R channel, was used to quantify the amount of protein retained on the bead surfaces for each of the 8 demultiplexed beads, as described in Suppl. Figure 15C.

#### C.5 Data Analysis

The mean fluorescence response in the Red-R channel for each beads color was calculated. To account for background signals, the responses from the no-expression control were subtracted from all measurements. Each specific target's response was then normalized to the response of Protein A beads coated with anti-SNAP antibody, as described in Suppl. Figure 15D.

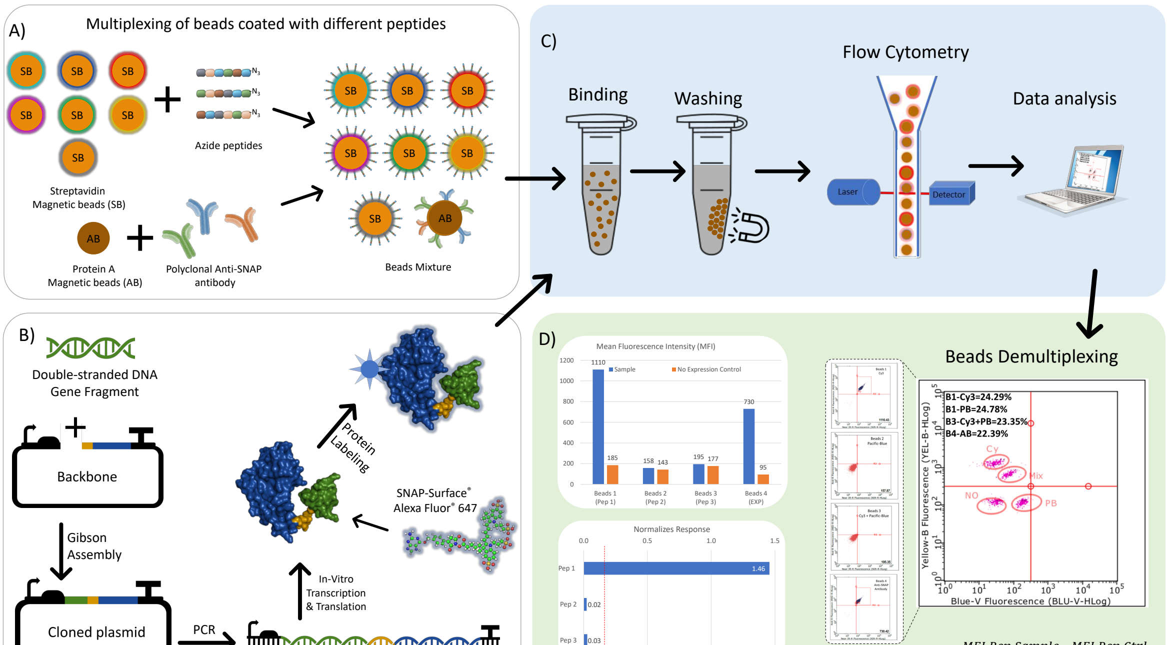


$$Normalizes\ Response = \frac{MFI\ Pep\ Sample - MFI\ Pep\ Ctrl}{MFI\ Exp\ Sample - MFI\ Exp\ Ctrl}$$

Suppl. Figure 15: Schematic overview of the experimental workflow (A) Magnetic beads labeling with various targets. Streptavidin-coated magnetic beads (SB) are incubated with color coded Biotin-DNA-DNCO linkers then azide modified peptides. Simultaneously, Protein A magnetic beads (AB) are used for binding with polyclonal anti-SNAP antibodies. Following this, all beads are mixed and ready for the binding step. (B) Schematic representation of the gene cloning and protein labeling process. Double-stranded DNA gene fragments are assembled into a plasmid backbone using Gibson Assembly, followed by PCR amplification. The PCR product is then used for in vitro transcription and translation to produce the SNAP-tagged protein, which is subsequently labeled with the SNAP-Surface® Alexa Fluor® 647 fluorescent dye. (C) Binding of expressed protein to the various beads. Following the binding and washing steps, the bead samples are passed through a flow cytometer. Here, the beads are excited by a laser, and the emitted fluorescence is detected by the instrument. The data is then processed for analysis. (D) Bead demultiplexing and data analysis. The mean fluorescence intensity (MFI) for each bead population is shown, with comparisons between samples and no-expression controls. Fluorescence data are normalized to correct for background signals. A representative flow cytometry plot shows bead populations based on fluorescence profiles, with distinct clusters identified for different bead colors. The MFI values for samples are shown alongside control responses, providing a clear indication of the relative fluorescence intensities.

### D Ancestral domain reconstruction on families of homologous proteins

#### D.1 Material and Methods

Within WW domains from PFAM seed (PF00397), twenty-four domains fell within or near the cluster associated to type IV domains. Complete protein sequences were retrieved and we conducted two searches for distant homologs using HMMER ([91], global homology search against the alphafold-uniprot50 database) and pLM-Blast ([92], local homology search with a 15-amino acid window length against the ECOD50 database). Subsequently, the protein families were enriched by NCBI pblast against the clustered NR database to obtain a number between 200 and 600 proteins per family.

Families were aligned with MAFFT [93] starting from partial alignments obtained using HMMER and pLM-BLAST with the ≪ add ≫ option. Subsequently, the resulting alignments underwent cleaning by removing sequences that shared less than 50 homologous residues with the original PFAM query or exhibited significant evolutionary divergence (computed with the amino-acid mutation matrix LG and Gamma parameter discretized in four classes [89, 88]). Furthermore, fragments of aligned sequences shorter than 5 amino acids, aligned with fewer than 10 other proteins, or not shared with the query were excluded. In each instance, the potential WW domains within the sequences were retained, along with the flanking four residues on each side.

Tree reconstruction and ASR were performed using IQ-Tree in the same way, employing the best evolutionary model identified by ModelFinder [94] among protein models plus the Gamma parameter (mutation matrix LG and four classes Gamma parameter). Trees were rooted with sequences belonging to outgroups (split between prokaryotes/eukaryotes or between plants and metazoa/fungi).

#### D.2 Phylogeny reconstruction on proteins containing type IV domains

We conducted rigorous phylogenetic reconstructions employing proteins homologous to potential type IV domain-containing proteins extracted from the WW domains seed. The divergence between these homologous proteins is substantial, with a mean divergence rate within families of 67%, reaching up to 85%. Consequently, these proteins display a diverse range of WW domain subtypes, although most remain type IV domains, followed by type II/IIIdomains. This observation supports the hypothesis that the divergence of WW domains is a very ancient event, as they are shared by all known eukaryotes [52, 95], with a split date estimated between 1 and 2 billion years ago [96].

For several protein families (Q6CEL8, D0MTH1, D0N5C9, D0MTN7, see example in Suppl. Figure 18A.), despite being able to establish a remote homology between proteins containing different types of WW domains, the divergence was too high to reconstruct a continuous evolutionary path between the domains. This is due notably to the significant number of indel events that occurred leading to poor domain reconstruction, as evidenced by the scarce HMM alignments extended with MAFFT.

In contrast, for other protein families like Q75CN9, Q59KZ2, A3LXA7, A8P5N1, F1SC59, and A7SG38, see Suppl. Figure 18B. and C., we were able to establish a link between the evolution of different domains and their common ancestors. However, the homology between these families is so distant that the low number of events leading to domain diversification prevents us from pinpointing multiple ancestors along branches of the trees that lack splits. Consequently, we obtain long branches connecting domain IV and I, devoid of any intermediate sequences, which severely limits our understanding of the domains' evolution. In this scenario, RBM-sampled paths could prove beneficial in augmenting the information provided by conventional ASR, which is unable to reliably assess ancestors between sequences connected by very long branches for which the phylogenetic signal is saturated. On the contrary, RBMs can generate hypotheses regarding probable evolutionary paths, anchored at their endpoints by the most reliable ancestors.

Finally, a few protein families (B3RKR0, A8P5N1, B0DLC8, A7SG38, C4WY86, NP001351424, see Suppl. Figure 18D., E., F.) connect smoothly all subtypes of domains. In these cases, the ancestors are classified as type I, II, or IV, while the empty region of the graph remains devoid of sequences. Interestingly, the classification and distribution of ancestral sequences in this scenario aligns with the distribution of modern sequences. Notably, family B3RKR0 contains a protein (CAF1230585, an unnamed protein product from *Adineta ricciae*) containing a type I WW-domain that is close to the empty region of the graph. This unique protein is likley to have emerged from type IV WW domain-containing proteins, as confirmed by a blast of the full protein that reveals close homologous proteins containing type IV WW domains.

The reconstruction of ancestral domains by ASR, which attempts to connect highly divergent proteins, presents significant challenges. Despite these limitations, we successfully identified several homologous proteins for which a shared evolutionary history can be inferred. The RBM classification enabled us to predict the expected ligand affinities of hundreds of domains, both for contemporary uncharacterized protein screening and for putative sampled ancestral sequences. This approach addresses the usual constraint of ASR, which is that only a limited number of potential ancestral proteins can be tested experimentally [97, 29]. Our *in silico* approach revealed that, although rare, the transition between WW subtype domains occurred independently on multiple lineages (Suppl. Figure 18D., E., F.). In the 24 families studied, we confidently identified switches from type II/III to IV subtypes in 12 families, between type I and II/III in 10 families, and a potential switch between I and IV for only one family (Suppl. Figure 18F.). This demonstrates that, contrary to the expectations derived from the reconstruction obtained solely from seed WW-domain alignments, type IV domains are more likely to have derived from type II/III domains.

To evaluate the evolutionary reliability of the tree constructed solely on domain alignments, as was done in the seed analysis, we assessed the concordance between the trees constructed from full protein alignments and the positions of these domains once added to the seed-derived tree. The mean quartet distance between the two trees was 0.27 (ranging between 0.15 and 0.44), indicating that approximately 73% of quartets from one tree are concordant with quartets from the other tree. This result suggests that despite an overall agreement, incorporating additional positions (and consequently evidence) for the tree inference substantially alters the resulting topology. While the seed tree is highly informative from a fitness and evolutionary perspective, it does not reproduce phylogenetic relationships as observed in independent homologous proteins' families.

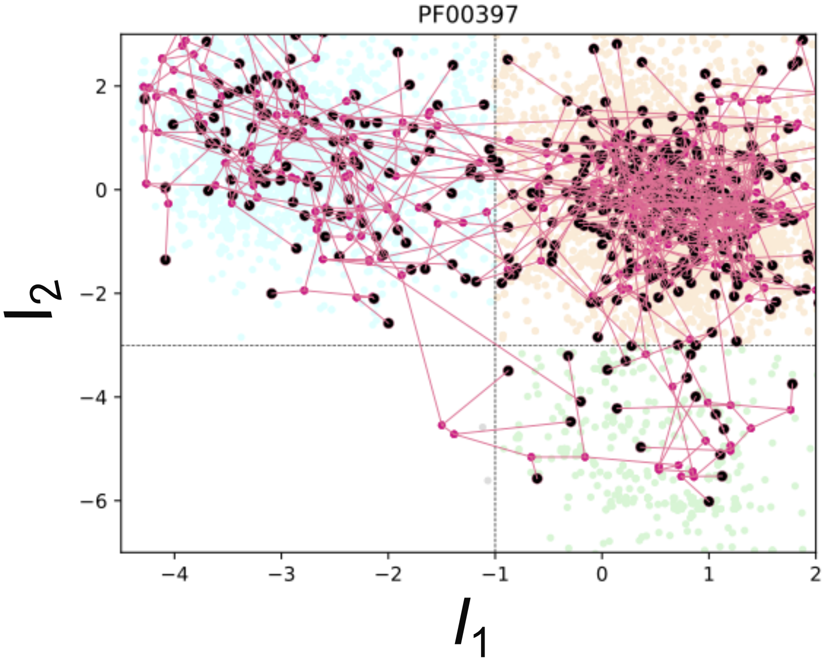


Suppl. Figure 16: **Paths derived from ancestral protein reconstruction.** 2D projections of the phylogenetic tree build from WW domain seed on the two specificity-related RBM hidden units. Black dots: modern sequences; Purple dots: average ancestral sequences sampled from the posterior distribution.

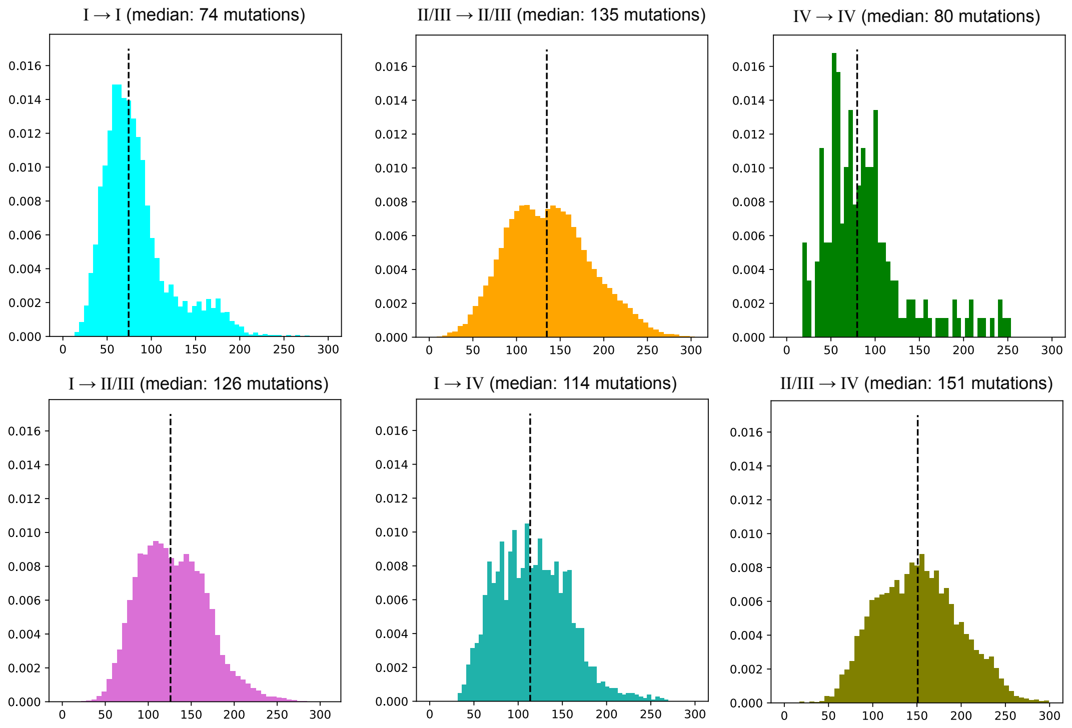


Suppl. Figure 17: **Path lengths obtained from PFAM WW-domain seed.** Distribution of paths lengths, expressed as expected number of mutations, between randomly chosen domains belonging to different subtypes.

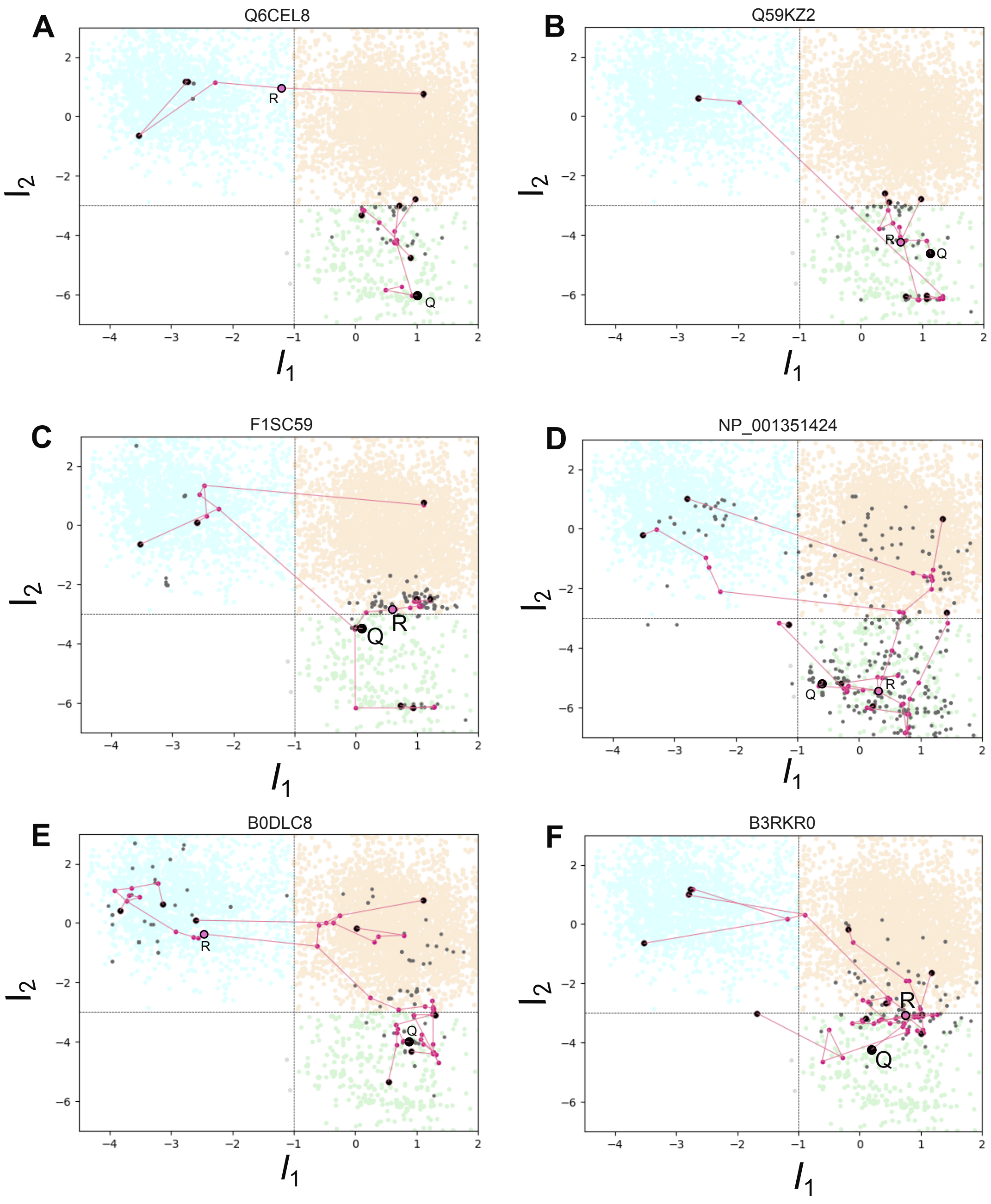


Suppl. Figure 18: **Paths derived from ancestral protein reconstruction.** 2D projection of the evolutionary path linking domains homolog to Q6CEL8 - Peptidyl-prolyl cis-trans isomerase from *Yarrowia lipolytica*, to Q59KZ2 - Peptidyl-prolyl cis-trans isomerase from *Candida albicans*, to F1SC59 - mRNA (2'-O-methyladenosine-N(6)-)-methyltransferase from *Sus scrofa*, to NP001351424 - Peptidyl-prolyl cis-trans isomerase from *Mus musculus*, to B0DLC8 - WW domain-containing protein from *Laccaria bicolor*, to B3RKR0 - Cap-specific mRNA (nucleoside-2'-O-)-methyltransferase 1 from *Trichoplax adhaerens*. These proteins are annotated Q on each panel and the root of the tree is R.
